# VGluT3 BNST neurons transmit GABA and restrict feeding without affecting rewarding or aversive processing

**DOI:** 10.1101/2025.01.01.631003

**Authors:** Annie Ly, Rachel Karnosky, Emily D. Prévost, Hayden Hotchkiss, Julianne Pelletier, Robert L. Spencer, Christopher P. Ford, David H. Root

## Abstract

The bed nucleus of the stria terminalis (BNST) is involved in feeding, reward, aversion, and anxiety-like behavior. We identify BNST neurons defined by the expression of vesicular glutamate transporter 3, VGluT3. VGluT3 neurons were localized to anteromedial BNST, were molecularly distinct from accumbal VGluT3 neurons, and co-express vesicular GABA transporter (VGaT). Cell-type specific presynaptic processes were identified in arcuate nucleus (ARC) and the paraventricular nucleus of the hypothalamus (PVN), regions critical for feeding and homeostatic regulation. Whole-cell patch-clamp electrophysiology revealed that, while these neurons co-express VGluT3 and VGaT, they functionally transmit GABA to both ARC and PVN, with rare glutamate co-transmission to ARC. Neuronal recordings of VGluT3 BNST neurons showed greater calcium-dependent signaling in response to sucrose consumption while sated compared with fasted. When fasted, optogenetic stimulation of BNST VGluT3 neurons decreased sucrose consumption using several stimulation conditions but not when stimulation occurred prior to sucrose access, suggesting that BNST VGluT3 activation concurrent with consumption in the fasted state reduces feeding. BNST VGluT3 activation during anxiety-like paradigms (novelty-suppressed feeding, open field, and elevated zero maze) and real-time place conditioning resulted in no changes in anxiety-like or reward/aversion behavior. We interpret these data such that VGluT3 BNST neurons represent a unique cellular population within the BNST that provides inhibitory input to hypothalamic regions to decrease feeding without affecting anxiety-like or reward/aversion behavior.

## INTRODUCTION

In the late 1970s, George Alheid embarked on what was intended to be a “brief excursion into neuroanatomy” before investigating the functionality of the extended amygdala(1). Over a half century later, the extended amygdala and its subdivisions and cell-types remain not wholly understood, as it continues to be discovered that its neuroanatomy and functionality are diverse, particularly the bed nucleus of the stria terminalis (BNST).

Historically, the BNST was subdivided according to medial-lateral, dorsal-ventral, and anterior-posterior axes based on tract tracing and immunohistochemical expression(2-13). The anterior BNST has dense projections to the paraventricular nucleus of the hypothalamus (PVN)(6, 9, 14-16). The anterior BNST can also be further subdivided into the anteromedial and anterolateral BNST. The anteromedial BNST has projections to the nucleus accumbens, arcuate nucleus (ARC), central amygdala, supramammillary and tuberomammillary nuclei, ventral tegmental area (VTA), and ventrolateral periaqueductal gray(6, 9, 16). The BNST, including the anteromedial BNST, has a strong role in feeding regulation(17-23), as well as in the expression of anxiety-like behavior(24-36).

With novel tools to probe the transcriptomics of neuroanatomically distinct brain regions in an unbiased fashion, it was discovered that the human and rodent BNST have a high level of transcriptional diversity(37, 38). Single-cell RNA sequencing data from mouse BNST revealed 41 transcriptionally distinct cell-types with 86% of cells expressing GABAergic genetic markers and the remaining 14% expressing glutamatergic genetic markers (38). Parcellating the behavioral functionality of the BNST is non-trivial, with several studies suggesting dichotomous, even opposing, roles in feeding, anxiety, and emotional valence depending on cell-type and circuit involvement(17, 39-42).

Here, we identified a cell-type within the anteromedial BNST that expressed the vesicular glutamate transporter type 3 (VGluT3). BNST VGluT3 neurons were distinct from striatal VGluT3 neurons due to the absence of the cholinergic marker choline acetyltransferase that classify striatal VGluT3-expressing cholinergic interneurons (43-45). Complexity of the VGluT3 cell-type within the anteromedial BNST was demonstrated using in situ hybridization and revealed that the majority of BNST neurons expressing VGluT3 mRNA co-expressed the vesicular GABA transporter (VGaT). The presynaptic vesicular protein synaptophysin from BNST VGluT3 neurons was identified locally within the BNST and also nucleus accumbens, PVN, ARC, and VTA. Additional immunolabeling and whole-cell patch-clamp electrophysiology experiments indicated that, while these neurons express VGluT3 protein, they functionally transmit GABA to PVN and ARC with rare glutamate co-transmission to ARC. Behaviorally, VGluT3 neurons showed greater calcium-related activity in response to sucrose consumption in the sated state compared with the fasted state, suggesting they may be involved in reducing sucrose intake. Optogenetic stimulation of BNST VGluT3 neurons resulted in decreased sucrose consumption in response to several stimulation conditions when sucrose was available but not when stimulation occurred prior to sucrose access. Optogenetic stimulation of VGluT3 neurons when mice were fasted resulted in no reliable changes in anxiety-like behavior. A real-time place preference test indicated that stimulation of VGluT3 BNST neurons is not inherently rewarding or aversive. Our results indicate an anteromedial BNST region that expresses a distinct genetic glutamatergic marker with dense inhibitory projections to hypothalamic regions. These VGluT3 BNST neurons are sensitive to internal state, such that its activation decreases feeding when fasted. VGluT3 activation had no effect on anxiety-like behavior, nor did it have significant emotional valence. Together, VGluT3 BNST neurons may represent a unique target for manipulating the neuronal regulation of food consumption or energy balance without affecting emotionally valent behaviors.

## MATERIALS AND METHODS

### Animals

VGluT3-IRES::Cre knock-in mice (4-5 months old; B6;129S-Slc17a8tm1.1(cre)Hze/J; Stock #028534), VGaT-IRES::Cre knock-in mice (4-5 months old; Slc32a1tm2(cre)Lowl/J; Stock #016962), VGaT-IRES::FlpO (B6.Cg-Slc32a1tm1.1(flpo)Hze/J, Stock #029591), and wildtype C57Bl6/J mice (Stock #000664) were purchased from The Jackson Laboratory (Bar Harbor, ME). Ai193 mice were received from The Allen Institute and crossed with double transgenic VGluT3::Cre/VGaT::Flp mice to create triple transgenic Ai193/VGluT3::Cre/VGaT::Flp mice. These mouse lines were bred and maintained at the University of Colorado Boulder and the University of Colorado Anschutz Medical Campus. Wildtype mice were used for the in situ hybridization experiment. All mice were group-housed by sex (5 mice/cage) under a reversed 12hr:12hr light/dark cycle (lights on at 22:00) with access to water ad libitum. All behavior experiments were performed during the dark phase of the light cycle. The experiments described were conducted in accordance with the regulations by the National Institutes of Health Guide for the Care and Use of Laboratory Animals and approved by the Institutional Animal Care and Use Committee at the University of Colorado Boulder as well as at the University of Colorado Anschutz Medical Campus.

### Surgery

Mice were anesthetized in a gasket-sealed induction chamber at 3% isoflurane gas. After confirming a surgical plane of anesthesia, isoflurane was continuously delivered at 1-2% concentration while the mouse was secured in the stereotactic instrument. For synaptophysin histology, VGluT3-IRES::Cre mice were unilaterally injected with AAV1-hSyn-FLEX-GFP-2A-Synaptophysin-mRuby (Stanford Vector Core, 5 × 1012 titer, 400 nL volume; 100 nl/min rate; +0.65 mm anteroposterior, ±0.6 mm mediolateral, −4.2 mm dorsoventral coordinates). For optogenetic stimulation experiments, AAV8-hSyn-DIO-CoChR-GFP (UNC Vectore Core) or AAV8-hSyn-DIO-GFP (Addgene) were injected bilaterally (5 × 1012 titer, 400 nL volume per hemisphere; 100 nl/min rate; +0.60 mm anteroposterior, ±0.6 mm mediolateral, −4.5 mm dorsoventral coordinates). For in vivo monitoring of calcium-dependent neural signaling, VGluT3-IRES::Cre mice were unilaterally injected with AAV1-hSyn-FLEX-GCaMP6m (Addgene) using the same surgical parameters as previously described. All injections were done using an UltraMicroPump, Nanofil syringes, and 35-gauge needles (Micro4; World Precision Instruments, Sarasota, FL). Syringes were left in place for 10 min following injections and slowly withdrawn. For optogenetics, bilateral fibers were implanted at a 10° angle (+0.60 mm anteroposterior, ±1.35 mm mediolateral, −4.3 mm dorsoventral coordinates). For fiber photometry, optic fibers (400-μm core diameter, 0.66 NA, Doric Lenses) were implanted slightly dorsal to the injection sites (+0.60 mm anteroposterior, ±0.6 mm mediolateral, −4.3 mm dorsoventral coordinates) and slowly lowered in discrete increments over the course of 25 minutes. All implants were secured with skull screws and dental cement. Mice were provided 3 days of postoperative care with 5 mg/kg carprofen (IP) and allowed 3-4 weeks of recovery before experimentation.

### Histology – Viral Verification and Synaptophysin

For viral verification, mice were anesthetized with isoflurane and perfused transcardially with 0.1M phosphate buffer followed by 4% (w/v) paraformaldehyde in 0.1M phosphate buffer, pH 7.3. Brains were extracted and cryoprotected in 18% sucrose solution in 0.1M phosphate buffer at 4° C overnight. Brains were cryosectioned to obtain coronal slices with BNST (30 μM). These coronal brain slices were mounted onto gelatin-coated slides and imaged for GFP fluorescent expression on a Zeiss widefield Axioscope. Any off-target or no GFP expression were excluded from analysis. For synaptophysin histology, slides were imaged on a Nikon A1R confocal (20X) with 3 μM steps z-stacked images at 512 × 512 pixel resolution. For each z-stacked image, the maximum intensity of each z-stack was measured and collapsed across all stacks using FIJI software. All reproductions of the mouse brain atlas for histology were derived from the Franklin and Paxinos atlas (46).

### Histology – In situ Hybridization

For in situ hybridization, C57BL6/J mice (8-12 weeks old) were anesthetized with isoflurane and perfused transcardially with RNase-free 0.1M phosphate buffer followed by 4% (w/v) paraformaldehyde in 0.1M phosphate buffer, pH 7.3. Brains were rapidly extracted and flash-frozen with isopentane. Brains were cryosectioned to obtain coronal slices (18 μM) from +1.10mm to +0.02mm bregma and were immediately mounted to Fisher SuperFrost Plus slides. Sections were then heat-treated and exposed to protease digestion followed by hybridization with target probes to mouse VGluT3 and VGaT mRNA (ACDBio). Sections were further treated with Opal dyes specific to each probe. Slides were coverslipped in Prolong Diamond with DAPI (ThermoFisher) and imaged on a Nikon A1R confocal (20X) with 3 μM steps z-stacked images at 512 × 512 pixel resolution. Images were further analyzed in FIJI, such that suitable regions of interest were defined as particles with an area over 10 μM2 and circularity. Particles that overlapped in expression for VGluT3 and VGaT were considered to be co-expressing VGluT3 and VGaT.

### Histology – Immunohistochemistry

In wildtype mice, frozen brain sections were thawed and washed 3 times for 10 minutes each with 0.1M phosphate buffer. Sections were then incubated in blocking buffer (4% bovine serum albumin and 0.3% TritonX-100 in 0.1M phosphate buffer, pH 7.3) for 1 hour at room temperature. Primary antibody solutions were rabbit anti-VGaT (Synaptic Systems #131003) and guinea pig anti-VGluT3 (Synaptic Systems #135204), diluted at 1:500 in blocking buffer solution. For Ai193/VGluT3::Cre/VGaT::Flp mice, the primary antibodies were goat anti-ChAT (Invitrogen PA518518), rabbit anti-tdTomato (Takara 632496), and mouse anti-GFP (Takara JL-8). Sections incubated in primary antibodies overnight at 4°C and then washed in 0.1M phosphate buffer (3×10 minutes). Secondary antibodies were donkey anti-rabbit Alexa594 (Jackson Immunoresearch #711585152) and donkey anti-guinea pig Alexa647 (Jackson Immunoresearch #706605148), diluted at 1:200 in blocking buffer. Sections incubated for 2 hours in antibody solutions and were washed again with 0.1 M PB (3×10 min). These sections were mounted onto subbed slides and coverslipped with ProLong diamond mounting medium. Slides were imaged with a NikonA1R confocal microscope (20X air objective and 100X oil objective) for VGluT3 and VGaT expression. Gain and power for each channel remained the same across images. Images were taken in a Z-stack consisting of 0.2μm steps. In post-processing, masks were generated for both the VGluT3 and VGaT layers using the Otsu method, and these binary matrices were used to remove any labels that did not overlap between the two layers, in order to calculate co-localization.

### Whole-Cell Patch-Clamp Electrophysiology

For slice preparation, mice were anesthetized with isoflurane and transcardially perfused with ice-cold cutting solution containing (in mM): 75 NaCl, 2.5 KCl, 6 MgCl2, 0.1 CaCl2, 1.2 NaH2PO4, 25 NaHCO3, 2.5 D-glucose, 50 sucrose. Coronal slices (240 μm) with the BNST, ARC, and PVN were cut in the same cutting solution that was used for transcardial perfusion. Slices were maintained at 32°C in aCSF containing (in mM): 126 NaCl, 2.5 KCl, 1.2 MgCl2, 1.2 NaH2PO4, 2.5 CaCl2, 21.4 NAHCO3, 11.1 D-glucose and 10 μm MK-801. After at least 30 minutes of incubation, slices were transferred to a recording chamber and continually perfused with 34 ± 2°C aCSF at a rate of 2 ml/min. All solutions were always bubbled with 95% O2, 5% CO2. When appropriate, picrotoxin (100 μM) was either bath applied or in the aCSF. DNQX (10 μM) was bath applied when appropriate.

All whole-cell recordings were performed using an Axopatch 200B amplifier (Molecular Devices). Data were acquired using an ITC-18 interface (Instrutech) and Axograph X software (Axograph Scientific) at 10 KHz and filtered to 2 KHz. Neurons were visualized on a BX51WI microscope (Olympus) with a LED and filter cube (ThorLabs). Samples that showed GFP fluorescence both at BNST cell bodies and fiber expression in the ARC and PVN were selectively used for whole cell recordings. Samples that did not show GFP fluorescence at BNST cell bodies nor fiber expression at the ARC or PVN were excluded. Widefield activation of CoChR was achieved with collimated light from a LED (470 nm) through the 40x water immersion objective. For whole-cell voltage-clamp recordings, cells were held at a voltage of 0mV and either −50mV or −60mV for the ARC and PVN, respectively. For the ARC, only cells with a series resistance less than 30 MΩ and an input resistance of more than 1 GΩ were recorded, as previously reported(47, 48). For the PVN, only cells with a series resistance less than 20 MΩ and an input resistance of at least 500 MΩ were recorded, as previously reported(49). Episode intervals were 30s with a single pulse of a 1ms width occurred at 2500ms within a 3000ms timebase. For CoChR validation, BNST cells were current-clamped, and a single pulse with a 1ms width occurred at 300ms in a 500ms timebase with episode intervals of 30s. Patch pipettes (2.5-3 MΩ) were pulled from borosilicate glass (World Precision Instruments). The internal pipette solution for all voltage-clamp experiments contained (in mM): 150 CsMeSO4, 10 HEPES (K), 0.1 EGTA, 0.1 CaCl2, 2 MgCl2, 0.2 Na2-GTP, 2 Na2-ATP, 4.6 Na-phosphocreatine, and 5 QX-314 bromide (pH 7.3, 280 mOsm). The internal pipette solution for current-clamp experiments contained (in mM): 135 KCl, 10 HEPES (K), 0.1 EGTA, 0.1 CaCl2, 2 MgCl2, 0.2 Na2-GTP, 2 Na2-ATP, and 4.6 Na-phosphocreatine (pH 7.3, 280 mOsm). All data processing occurred using Axograph (1.7.6). A baseline subtraction was applied based on a 100 ms range prior to the event. A sharp low-pass filter was applied with a cutoff frequency of 2000Hz. The maximum peak was detected after the stimulation event.

### Calcium Fiber Photometry Recordings

GCaMP6m was excited at two wavelengths (465nm and 405nm isosbestic control) with amplitude-modulated signals from two light-emitting diodes reflected off dichroic mirrors and then coupled into an optic fiber (50-52). The GCaMP signal and the isosbestic control signal were returned through the same optic fiber and acquired using a femtowatt photoreceiver (Newport, Irvine, CA), digitized at 1kHz, and then recorded by a real-time signal processor (Tucker Davis Technologies). For analysis of calcium fiber photometry recordings, custom-written MATLAB scripts were used and are available at www.root-lab.org/code. The isosbestic signal (405nm) and the GCaMP signal (465nm) were downsampled (10x) and peri-event time histograms were created surrounding events of interest (e.g., sucrose consumption, shock). For each trial, data were detrended by regressing the 405nm signal on the 465nm signal. The generated linear model was used to create a predicted 405nm signal that was subtracted from the 465nm signal to remove movement, photo-bleaching, and fiber bending artifacts (50). Baseline normalized maximum z-scores were taken from −10 to 0.01 seconds prior to event onset.

### Fiber Photometry - Sucrose Consumption

Mice were recorded in response to consuming 8% sucrose either in the sated state (free fed) or fasted state (fasted to 85-90% body weight). Mice were trained to consume within a 10-minute time period for 5 days before recordings. The apparatus (Med-Associates) was outfitted with a spout for a 250ml sipper bottle (Allentown, LLC). Mice were habituated to the fiber photometry cable at least 48 hours before recordings. The order of recordings during either the fasted or sated state were counter-balanced. A lick bout was defined as the occurrence of at least 2 licks. Onset of a lick bout was defined as the first lick with at least a 3 second prior interval in which no licks occurred.

### Fiber Photometry - Shock Exposure

Mice were recorded in response to five electric footshocks while in the fasted or sated state, the order of which was counter-balanced. In the fasted state, mice were limited to 85-90% of their original body weight, while they were given food ad libitum in the sated state. Mice were administered 0.5mA 500ms electric foot shocks in a novel environment with an average inter-trial interval of 60s.

### Fiber Photometry – Pavlovian Reward Conditioning

Mice were fasted to 85-90% body weight and trained to consume 8% sucrose within a 100-minute time period in a Pavlovian conditioning task. The apparatus (Med-Associates) was outfitted with a custom 3D-printed reward magazine, which can be found on www.root-lab.org/code. A single-speed syringe pump (Med-Associates) was used to deliver 30µl sucrose to the reward magazine. The task consisted of 40 presentations of a 10s conditioned stimulus (CS+) that co-terminated with sucrose reward delivery, which was 1.5s long. The task also included 40 presentations of a 10s conditioned stimulus (CS-) that had no programmable changes. The average inter-trial interval was 90s. Cue onset was defined as the time CS+ or CS-started sounding. Reward retrieval onset was defined as the time mice entered the magazine during cue presentation.

### Optogenetic Stimulation – Sucrose Consumption

Mice were fasted to 85-90% of their original body weight and trained to consume 8% sucrose within a 40-minute time period for 7 days before stimulation experiments began. The apparatus (Med-Associates) was outfitted with a spout for a 250ml sipper bottle (Allentown, LLC). Mice were habituated to the fiber optic cable at least 48 hours before stimulation experiments began. All stimulation conditions had a pulse width of 5ms and a frequency of 20Hz. Light intensity at the end of the tip was set to be approximately 10 mW. Stimulation conditions were determined based on previously published research(53-55). For open-loop stimulation, stimulation occurred in two 10-minute time blocks within the 40-minute time period, such that there was no stimulation in the first 10 minutes and from 20 to 30 minutes. For the calculated sucrose ratio, licks that occurred during the 10-minute blocks of stimulation were divided by the number of licks during the non-stimulation 10-minute blocks. For closed-loop stimulation, a stimulation train was only triggered by 3 licks with an inter-lick interval less than 1s. For prior stimulation, stimulation occurred continuously for 20 minutes in a novel environment before being placed in the behavior apparatus for sucrose consumption. For tonic stimulation, stimulation occurred continuously throughout the 40-minute session. At least 24 hours of no stimulation separated each stimulation condition. For closed-loop stimulation, prior stimulation, and tonic stimulation, licks were divided by licks from the preceding day when no stimulation occurred.

### Novelty-Suppressed Feeding

Mice were food-deprived for 24 hours before the start of the novelty-suppressed feeding task. For the novelty-suppressed feeding task, mice were placed in an open field box (40cm length x 40cm width x 35cm height, Stoelting) in which the center was illuminated at 40 lux. The center of the box contained a petri-dish weighed down by batteries. A standard food biscuit (approximately 5g) was attached to the petri-dish using a rubber band, in order to prevent mice from pulling the food biscuit away from the center. Photo-stimulation immediately began when mice were placed into the open field and lasted for 10 minutes. Light intensity at the end of the tip was set to be approximately 10 mW. Photo-stimulation was performed at 20 Hz with 5 ms pulse width using blue laser light (473 nm, CNI Laser). The center of the box was defined as 5% of the total area, while the periphery was defined as 50% of the total area and was limited to the edges of the box. Food approach latency was measured as the time it took the mice to approach the center with food. Novelty-suppressed feeding behavior was video recorded and quantified using ANY-Maze software.

### Open Field

Mice were fasted to 85-90% of their original body weight before being placed in an open field box (40cm length x 40cm width x 35cm height, Stoelting). Photo-stimulation immediately began when mice were in the box and lasted for 20 minutes. Photo-stimulation parameters were the same as previously described. The center of the box was defined as 5% of the total area, while the periphery was defined as 50% of the total area and was limited to the edges of the box. Approach latency was defined as the time it took for the center of the mice to reach the center of the box. Open field behavior was video recorded and quantified using ANY-Maze software.

### Elevated Zero Maze

Mice were fasted to 85-90% of their original body weight before being placed on an elevated zero maze apparatus (78cm height, 40cm diameter, 5cm track width, 30cm wall height, 31cm closed arm length, Accuscan Instruments). The apparatus was adjusted such that the original black circular track was covered with white Plexiglas for better video tracking, and the clear walls of the closed arms were taped with black light-blocking tape. Mice were placed in the center of one of the closed arms, and photo-stimulation immediately began. Photo-stimulation lasted for 10 minutes, and parameters were the same as previously described. Enter latency was defined as the time it took for the mice to reach the other closed arm, in which all four paws were in the other closed arm. Elevated zero maze behavior was video recorded and quantified using ANY-Maze software.

### Real-Time Place Preference

The real-time place preference box (40cm length x 40cm width x 35cm height, Stoelting) consists of a stripe chamber (18cm × 20cm), solid chamber (18cm × 20cm), and a connecting antechamber. The solid chamber had been modified with a peel-and-stick window film to prevent reflection. Mice were acclimated to the tethers during baseline real-time place preference with no photo-stimulation and with ad libitum food access. The next day, mice were subjected to 15-minute photo-stimulation on both the solid and stripe sides with the sides counter-balanced between mice. Photo-stimulation parameters were the same as previously described. The same experiment was repeated when mice were fasted to 85-90% of baseline body weight. Real-time place preference behavior was video recorded and quantified using ANY-Maze software.

### Statistical Analysis

Values are reported as mean ± standard error with bars depicted as shaded areas in fiber photometry traces. All statistical tests were performed with R (4.0.5). Results were subject to a two-way ANOVA analysis to test the effect of holding voltage (−50mV, 0mV), drug (baseline, picrotoxin), group (GFP, CoChR), and state (fasted, sated), where appropriate. Comparisons were made by paired or non-paired, two-tailed t-test with Bonferonni corrections.

## RESULTS

### Co-localization of vesicular markers of glutamate and GABA within the anteromedial BNST

To identify glutamatergic and GABAergic BNST neurons, we used RNAscope in situ hybridization to label VGluT3 and VGaT mRNA in wildtype mice (N=4, 2 female, 2 male). Expression of VGaT mRNA was abundant throughout BNST while VGluT3 mRNA expression appeared confined to the anteromedial BNST from AP +0.62 mm to AP +0.38 mm (**Figure 1A**). To test whether BNST VGluT3 neurons were distinct from striatal VGluT3 neurons expressed in cholinergic interneurons (44, 56), we generated triple transgenic Ai193/VGluT3::Cre/VGaT::Flp mice where VGluT3-Cre neurons expressed GFP and VGaT-Flp neurons expressed tdTomato. We found that BNST VGluT3 neurons lacked ChAT immunolabeling while striatal VGluT3 neurons co-expressed ChAT (Figure 1B), demonstrating that BNST VGluT3 neurons belong to a different class of neurons than striatal VGluT3 cholinergic interneurons.

**FIGURE 1.**
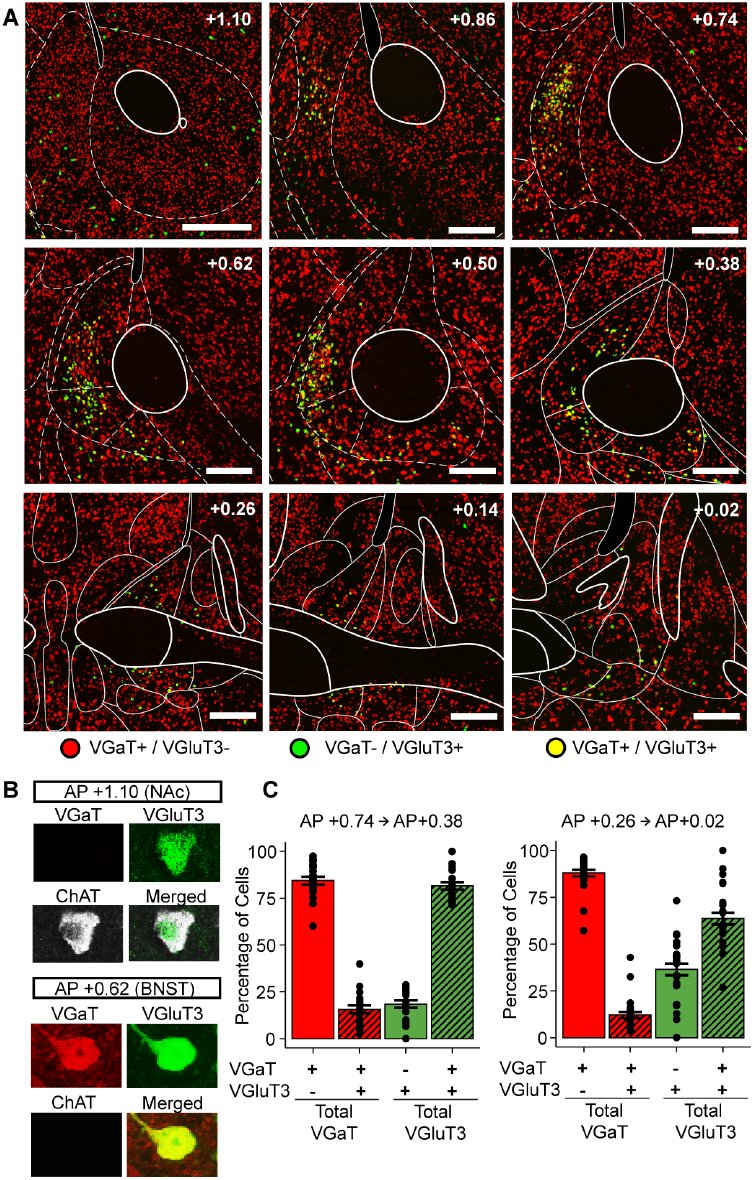
Co-expression of vesicular markers of glutamate and GABA within the anteromedial BNST. (A) Example RNAscope histology from AP +1.10 mm to AP +0.02 mm with VGaT mRNA in red and VGluT3 mRNA in green. N=4, 2 female, 2 male. Scale bar is 200 μm. (B) Nucleus accumbens (NAc) VGluT3 neurons express ChAT while BNST VGluT3 neurons co-express VGaT and lack ChAT. Magnification is 40X. (C) Quantification of VGluT3 and VGaT mRNA co-expression across AP axis of the BNST. Outlines are based on the Franklin and Paxino mouse brain atlas.

We next quantified the co-expression of VGaT and VGluT3 mRNA in BNST between AP +0.74 mm to AP +0.38 mm (anterior) and between AP +0.26 mm to AP +0.02 mm (posterior; N=4 mice, approximately 20 sections/mouse) (**Figure 1C**). Anteriorly, most VGluT3-expressing neurons co-expressed VGaT (81.5% ± 1.95) and the VGluT3-expressing neurons were a small subpopulation out of the total VGaT-expressing population (15.6% ± 2.13). There were fewer VGluT3-only neurons (18.5% ± 1.95) anteriorly compared with VGaT-only neurons (84.4% ± 2.13). Posteriorly, most VGluT3-expressing neurons still co-expressed VGaT but were less than the anterior portion (63.6% ± 3.12), and the percentage of VGluT3-only neurons increased to 36.4% ± 3.12. Like anterior coordinates, most BNST neurons were VGaT-only (87.9% ± 1.67) and the percent of VGluT3-expressing neurons was low out of the total VGaT-expressing population (12.1% ± 1.67).

### VGluT3 neurons in the anteromedial BNST form synaptic projections with hypothalamic brain regions

After identifying the unique transcriptional and anatomical profile of VGluT3 BNST neurons, we next aimed to determine their efferent circuitry. Using AAV1-hSyn-FLEX-GFP-2a-Synaptophysin-mRuby that labels cell bodies and axons within GFP and the presynaptic vesicular protein synaptophysin within mRuby (57), we identified the vesicular synaptophysin distribution of VGluT3 neurons (N=6, 3 female, 3 male) (**Figure 2A**). Discrete sites of pre-synaptic vesicular expression were identified in the nucleus accumbens shell (AP +0.74), anterior BNST (AP +0.26), posterior BNST (AP −0.10), paraventricular hypothalamus (AP −0.94), arcuate nucleus (AP −1.58), and the ventral tegmental area (AP −−−−3.16) (**Figure 2B**). The densest mRuby expression that was not within the BNST itself was the arcuate nucleus (ARC), followed by the paraventricular hypothalamus (PVN). Negligible to no expression was observed elsewhere, including lateral habenula that receives co-transmitted glutamate and GABA from VTA (58). In order to validate that genetic expression of VGluT3 also resulted in protein expression, we labeled VGluT3 and VGaT protein in the primary targets of BNST VGluT3 neurons, the PVN (**Figure 2C**) and ARC (**Figure 2D**) (N=8, 4 female, 4 male). We found that approximately half of VGluT3 protein was co-localized with VGaT protein in each region (50.1% ± 0.01% in the PVN and 50.3% ± 0.02% in the ARC) (**Figure 2E**).

**FIGURE 2.**
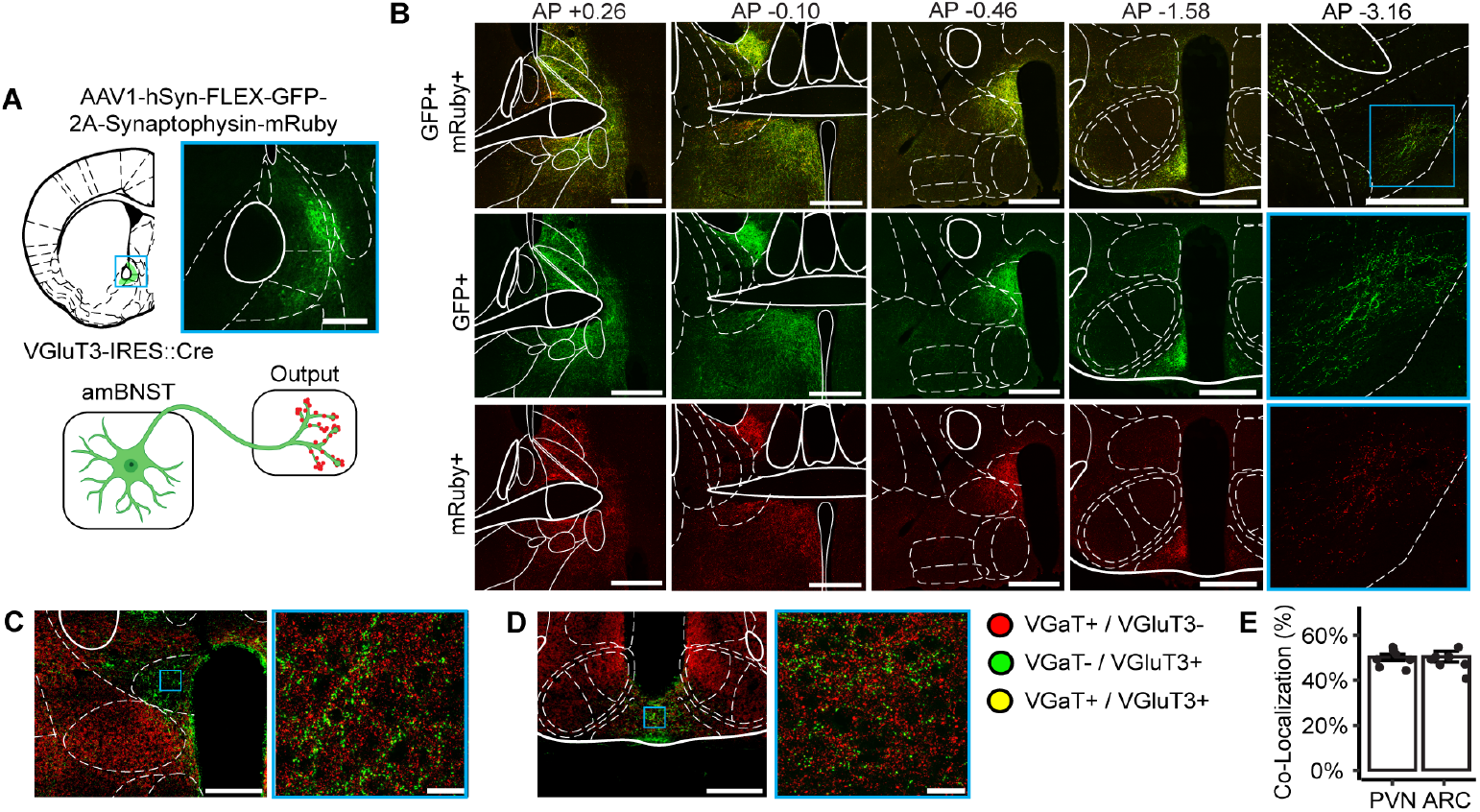
VGluT3 BNST neurons primarily synapse within the BNST and with hypothalamic brain regions. (A) VGluT3-IRES::Cre mice were injected with FLEX-GFP-2A-Synaptophysin-mRuby, labeling cell bodies and fibers in GFP and synaptophysin-expressing vesicles in mRuby. Scale bar is 500 μm. N=6, 3 female, 3 male. (B) GFP and mRuby expression throughout the BNST, paraventricular nucleus of the hypothalamus (PVN), arcuate nucleus (ARC), and the ventral tegmental area (VTA). Scale bar is 500 μm. (C) VGluT3 and VGaT protein labeling in the PVN with VGluT3 in green and VGaT in red. Scale bar is 200 μm. High magnification image scale bar = 20 μm. (D) VGluT3 and VGaT protein labeling in the ARC. Scale bar is 500 μm. High magnifcation image scale = 20 μm. (E) Quantification of VGaT co-localization out of total VGluT3 protein in the PVN and ARC. Images outlined in blue in (B), (C), and (D) are 100x zoomed in images.

### VGluT3 BNST neurons, while genetically expressing the capability to transmit glutamate, functionally transmit GABA and rarely transmit glutamate to the ARC

The co-expression of VGluT3 and VGaT mRNA in BNST neurons and in the projection targets of these neurons suggests BNST VGluT3 neurons may co-transmit glutamate and GABA. In order to investigate the synaptic functionality of VGluT3 and VGaT co-expression, VGluT3-IRES::Cre mice were injected with the Cre-dependent, high-photocurrent excitatory opsin CoChR (59, 60) and BNST VGluT3 axons were optogenetically stimulated while performing whole-cell recordings in the ARC (**Figure 3B**). This optogenetic strategy to stimulate action potentials from BNST VGluT3 neurons was validated ex vivo (**Figure 3A**). The experimental design to obtain both a GABA-mediated inhibitory post-synaptic current and an AMPA-mediated excitatory post-synaptic current (EPSC) from the same cell necessitated an accurate reading of the chloride reversal potential. Thus, the chloride reversal potential was experimentally determined in the ARC while washing on the AMPA receptor antagonist, DNQX, which was −50 mV (**Figure 3C**). In order to experimentally determine if VGluT3 BNST neurons transmit both GABA and glutamate, ARC neurons were voltage-clamped at both 0 mV and −50 mV while the terminals were optogenetically stimulated. Example traces of an ARC neuron showed an optogenetic post-synaptic current at 0 mV, which was abolished following application of the GABAA receptor antagonist, picrotoxin; no post-synaptic current was observed under baseline conditions at −50 mV (**Figure 3D**). In the ARC, there was a significant effect of drug and voltage (F(1,24)=19.1, p= 0.0002) (N=7 cells, 2 female, 2 male) (**Figure 3E**). Post-hoc analysis showed that at baseline conditions, there was a significant difference in amplitude between −50 mV and 0 mV (t(6.14) = −4.39, p = 0.004, −50 mV = −6.12 ± 17.25, 0 mV= 260.59 ± 59.67). At 0 mV, there was a significant difference in amplitude between baseline and picrotoxin wash (t(6.01)=4.34, p=0.004, picrotoxin = −1.54 ± 3.89). Thus, picrotoxin significantly reduced the optogenetic post-synaptic current at 0 MV with no changes to amplitude at −50 mV. Next, in the continued presence of picrotoxin in the aCSF, we investigated if any recorded ARC neurons demonstrated a DNQX-sensitive post-synaptic current (N=16 cells, 2 female, 2 male). Only 6.525% of recorded neurons exhibited an optogenetic post-synaptic current at −50 mV, which was abolished with DNQX (**Figure 3F**). In order to experimentally validate GABA transmission, VGaT-IRES::Cre were similarly injected with CoChR and whole-cell recordings in the ARC were performed (**Figure 3H**). The optogenetic strategy to stimulate action potentials from VGaT-expressing neurons was validated ex vivo (**Figure 3G**). Similar to VGluT3::Cre projections, example traces from a single ARC neuron illustrated an optogenetic post-synaptic current at 0 mV, which was abolished following application of the GABAA receptor antagonist, picrotoxin; again, no post-synaptic current was observed under baseline conditions at −50 mV (**Figure 3I**). In the ARC, with VGaT BNST terminal stimulation, there was a significant effect of drug and voltage (F(1, 36)=17.766, p=0.0001) (N=10 cells, 2 female, 3 male) (**Figure 3J**). Post-hoc analysis showed that at baseline conditions, there was a significant difference in amplitude between −50 mV and 0 mV (t(9.03)=-4.45, p=0.001, −50 mV = −16.4 ± 23.1, 0 mV= 805.6 ± 559.7). As expected, at 0 mV, there was a significant difference in amplitude between baseline and picrotoxin wash (t(9.07)=4.31, p=0.001, picrotoxin = 40.08 ± 35.4). In the continued presence of picrotoxin in the aCSF, we investigated if any recorded ARC neurons demonstrated a DNQX-sensitive post-synaptic current that could be induced by input from VGaT BNST neurons (N=14 cells, 2 female, 2 male). Only 7.14% of recorded neurons exhibited an optogenetic post-synaptic current at −50 mV, which was abolished with DNQX (**Figure 3K**). Thus, only a small minority of VGluT3 terminals functionally transmit glutamate.

**FIGURE 3.**
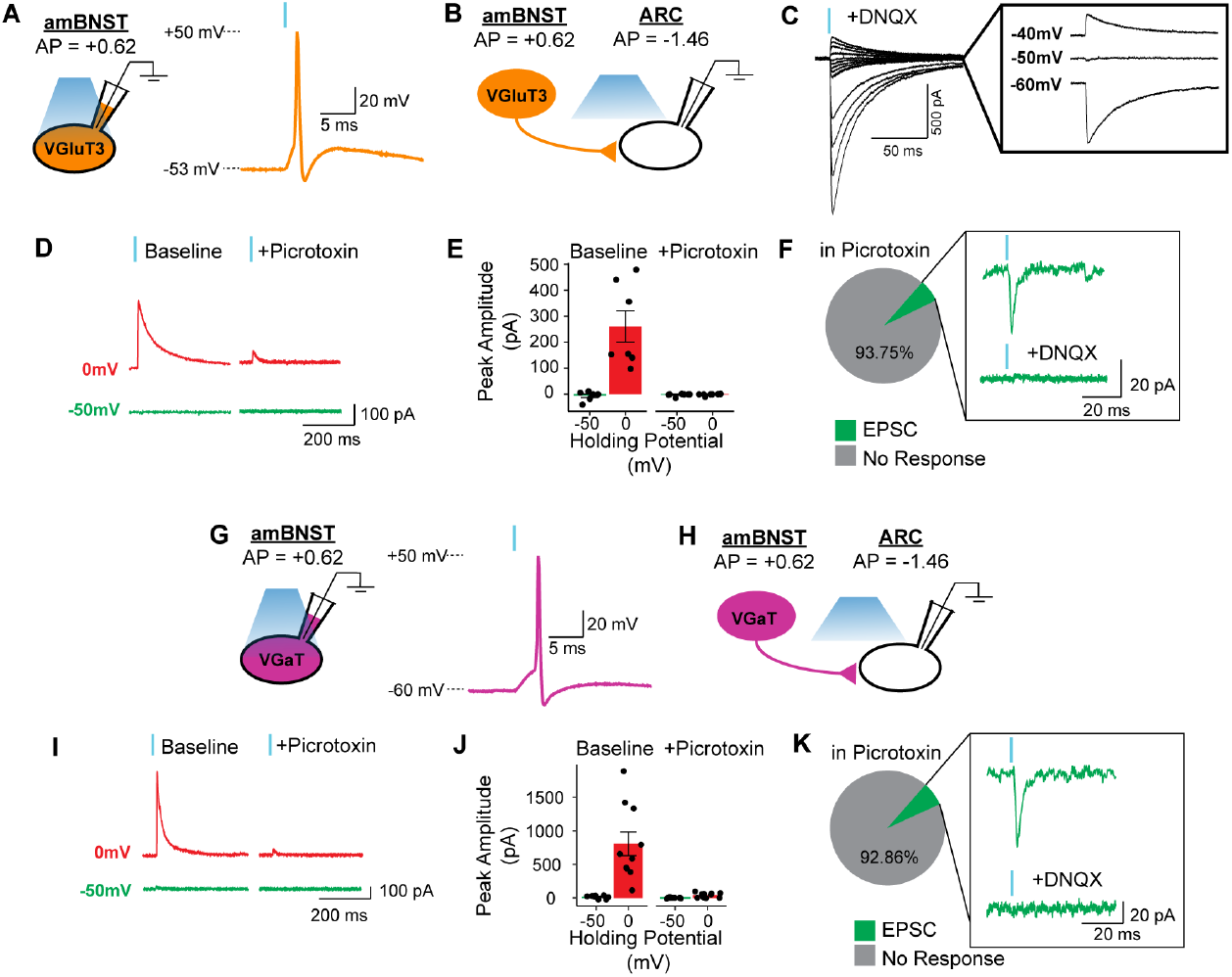
VGluT3 BNST neurons functionally transmit GABA and rarely transmit glutamate to the ARC. (A) Validation of optogenetic-induced action potential of a BNST VGluT3 neuron. (B) VGluT3 terminals were optogenetically stimulated while patching onto an ARC neuron. (C) Chloride reversal potential was experimentally determined of ARC neurons while washing on AMPA receptor antagonist, DNQX, which was −50 mV. (D) VGluT3 to ARC example traces of optogenetic post-synaptic currents under 0 mV and −50 mV at baseline and with GABAA receptor antagonist, picrotoxin, wash. (E) Picrotoxin wash application on VGluT3 optogenetic post-synaptic current at −50 mV and 0 mV. N=7 cells, 2 female, 2 male. (F) Percentage of recorded neurons that demonstrated a DNQX-sensitive post-synaptic current with picrotoxin in aCSF. N=16 cells, 2 female, 2 male. (G) VGaT terminals were optogenetically stimulated while patching onto an ARC neuron. (H) Validation of optogenetic-induced action potential of a VGaT-expressing neuron. (I) VGaT to ARC example traces of optogenetic post-synaptic currents under 0 mV and −50 mV at baseline and with GABAA receptor antagonist, picrotoxin, wash. (J) Picrotoxin wash application on VGaT optogenetic post-synaptic current at −50 mV and 0 mV. N=10, 2 female, 3 male. (K) Percentage of recorded neurons that demonstrated a DNQX-sensitive post-synaptic current with picrotoxin in aCSF. N=14 cells, 2 female, 2 male.

### VGluT3 BNST neurons functionally transmit GABA but not glutamate to the PVN

The second major output of VGluT3 BNST neurons was the PVN. We investigated if transmission from VGluT3 BNST neurons was different between the ARC and the PVN. VGluT3-IRES::Cre mice were injected with CoChR and whole-cell recordings in the PVN was performed (**Figure 4A**). Due to the variability in chloride reversal potential between cell-types and the necessity of an accurate reading, the chloride reversal potential was experimentally determined in the PVN while washing on the AMPA receptor antagonist, DNQX, which was −60 mV (**Figure 4B**). In order to experimentally determine if VGluT3 BNST neurons transmit both GABA and glutamate, PVN neurons were voltage-clamped at both 0 mV and −60 mV while the terminals were optogenetically stimulated. Example traces of a PVN neuron showed an optogenetic post-synaptic current at 0 mV, which was abolished following application of the GABAA receptor antagonist, picrotoxin; no post-synaptic current was observed under baseline conditions at −50 mV (**Figure 4C**). In the PVN, there was a significant effect of drug and voltage (F(1, 28)=16.1, p=0.0003) (N=8 cells, 2 female, 2 male) (**Figure 4D**). Post-hoc analysis showed that at baseline conditions, there was a significant difference in amplitude between −60 mV and 0 mV (t(7.24)=-4.29, p=0.003, −60 mV = −72.47

**FIGURE 4.**
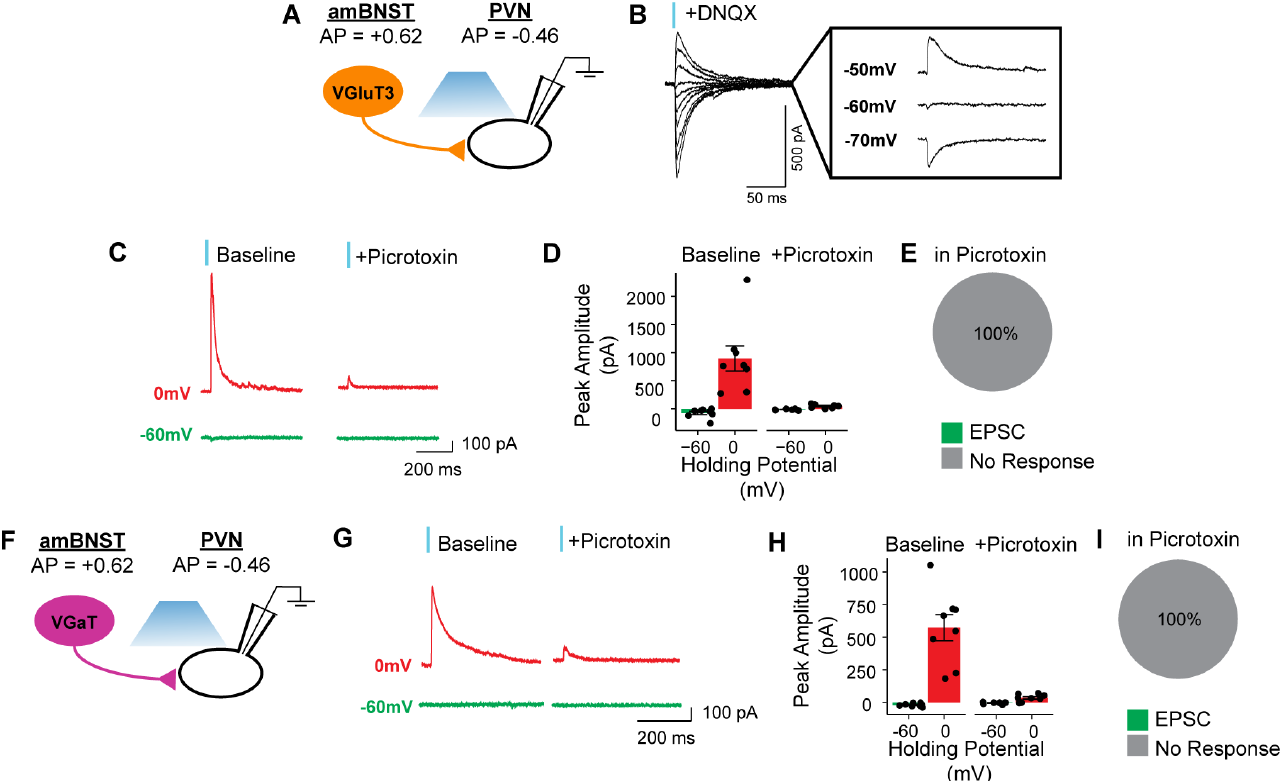
VGluT3 BNST neurons functionally transmit GABA but not glutamate to the PVN. (A) VGluT3 terminals were optogenetically stimulated while patching onto a PVN neuron. (B) Chloride reversal potential was experimentally determined of PVN neurons, which was −60 mV. (C) PVN example traces of VGluT3 optogenetic post-synaptic currents under 0 mV and −60 mV at baseline and with picrotoxin wash. (D) Picrotoxin wash application on VGluT3 optogenetic post-synaptic current. N=8 cells, 2 female, 2 male. (E) With picrotoxin in aCSF, no recorded neurons demonstrated an excitatory post-synaptic current. N=17 cells, 2 female, 2 male. (F) VGaT terminals were optogenetically stimulated while patching onto a neuron within the PVN. (G) PVN example traces of VGaT optogenetic post-synaptic currents under 0 mV and −60 mV at baseline and with picrotoxin wash. (H) Picrotoxin wash application on VGaT optogenetic post-synaptic current at −60 mV and 0 mV. N=8 cells, 2 female, 2 male. (I) With picrotoxin in aCSF, no recorded neurons demonstrated an excitatory post-synaptic current. N=22 cells, 2 female, 3 male.

± 84.29, 0 mV= 895.52 ± 631.28). At 0 mV, there was a significant difference in amplitude between baseline and picrotoxin wash (t(7.03)=3.79, p=0.006, picrotoxin = 47.99 ± 32.8). Next, in the continued presence of picrotoxin in the aCSF, we investigated if any recorded PVN neurons demonstrated a DNQX-sensitive post-synaptic current (N=17 cells, 2 female, 2 male). Compared to the ARC, no recorded neurons exhibited an optogenetic post-synaptic current at −60 mV (**Figure 4E**). VGaT-IRES::Cre were injected with CoChR while performing whole-cell recordings in the PVN (**Figure 4F**). Example traces from a single PVN neuron showed an optogenetic post-synaptic current at 0 mV, which was abolished following application of the GABAA receptor antagonist, picrotoxin; again, no post-synaptic current was observed under baseline conditions at −60 mV (**Figure 4G**). In the PVN, with VGaT BNST terminal stimulation, there was a significant effect of drug and voltage (F(1, 28)=30.2, p<0.00001) (N=8 cells, 2 female, 2 male) (**Figure 4H**). Post-hoc analysis showed that at baseline conditions, there was a significant difference in amplitude between −60 mV and 0 mV (t(7.03)=-5.91, p=0.0005, −60 mV = −17.86 ± 13.38, 0 mV= 573.26 ± 282.3). At 0 mV, there was a significant difference in amplitude between baseline and picrotoxin wash (t(7.13)=5.35, p=0.001, picrotoxin = 36.2 ± 27.6). In the continued presence of picrotoxin in the aCSF, no recorded neurons exhibited a DNQX-sensitive post-synaptic current at −60 mV with stimulation from VGaT BNST terminals (N=22 cells, 2 female, 3 male) (**Figure 4I**). Thus, different from the ARC, VGluT3 BNST neurons do not functionally transmit glutamate to the PVN.

### Distinct VGluT3 neural signaling to sucrose reward in a state-dependent manner

Both the ARC and the PVN are hypothalamic regions that regulate homeostatic state. The ARC exerts influence over food intake (61, 62) while the PVN is a critical node of the hypothalamic-pituitary-adrenal axis that also regulates food intake(63). Thus, we first investigated calcium-dependent signaling of VGluT3 BNST neurons in response to sucrose consumption and aversive footshock when mice were sated and fasted. VGluT3-IRES::Cre mice were injected with Cre-dependent GCaMP6m into the BNST and implanted with an optic fiber (**Figure 5A-B**). Mice were recorded in response to sucrose consumption when fasted or sated, the order of which was counter-balanced between mice (N=11, 6 female, 5 male) (**Figure 5C**). Sucrose consumption, as measured in licks, was significantly higher in the fasted state than the sated state (t(12.13)=-8.91, p<0.0001, sated = 115.8±29.9, fasted = 971.3±91.1) (**Figure 5D**). In spite of higher sucrose consumption in the fasted state, BNST VGluT3 neuronal activity was significantly higher when mice were sated compared with the fasted state (**Figure 5E-F**) (t(10)=4.06, p=0.002, sated = 9.91 ± 4.26, fasted = 5.17 ± 1.70). Mice were also recorded in response to footshock when fasted or sated, the order of which was counter-balanced between mice (N=6, 3 female, 3 male) (**Figure 5G**). In contrast to sucrose consumption, footshock increased VGluT3 neuronal activity in both the fasted and sated states (**Figure 5H-I**). We also assessed calcium-dependent VGluT3 activity during a Pavlovian conditioning task while mice were fasted to determine if reward learning had an impact on signaling. VGluT3 BNST neurons showed no signaling related to conditioned stimuli predicting sucrose delivery (CS+) or a stimulus predicting no sucrose delivery (CS-) (**Figure 5J-K**). When calcium-dependent signaling was time-locked to magazine entry to retrieve the sucrose reward during the cue, there was also no difference from baseline (**Figure 5L-M**). Overall, we interpret these results such that BNST VGluT3 neurons are sensitive to the interaction of palatable food reward and internal state, specifically when the subject is sated. However, BNST VGluT3 neurons are also sensitive to aversive stimuli that is not dependent on internal state.

**FIGURE 5.**
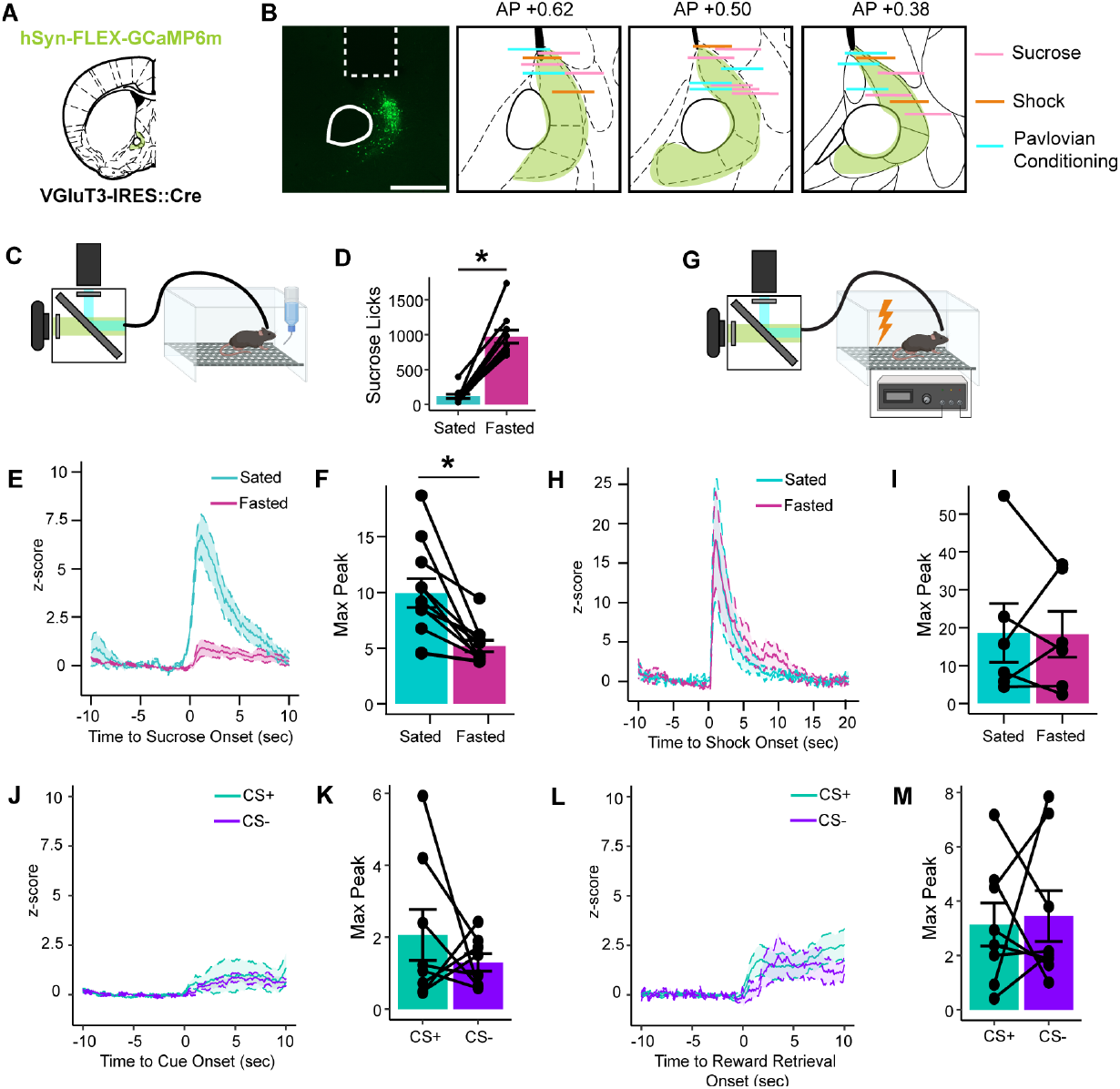
State-dependent BNST VGluT3 neuronal signaling of sucrose consumption. (A) VGluT3-IRES::Cre mice were injected with Cre-dependent GCaMP6m. Scale bar is 500 μm. (B) Histological verification of fiber placements. (C) While sated or fasted, mice were recorded in response to sucrose consumption. N=11, 6 female, 5 male. (D) Sucrose consumption in the sated and fasted states. (E) Averaged traces of VGluT3 calcium-dependent signaling in response to bouts of sucrose consumption between fasted and sated states.(F)Maximum peaks observed in response to sucrose of VGluT3 calcium-dependent signaling between the sated and fasted states. (G) While sated and fasted, mice were recorded in response to electric footshock. N=6, 3 female, 3 male. (H) Averaged traces of VGluT3 calcium-dependent signaling in response to electric footshock between fasted and sated states. (I) Maximum peaks observed in response to electric foot shock of VGluT3 calcium-dependent signaling between the sated and fasted states. (J) Averaged traces in response to a conditioned stimulus that predicted sucrose reward (CS+) and a neutral stimulus with no programmable effect (CS-). (K) Maximum peaks in response to CS+ and CS-. (L) Averaged traces of reward retrieval onset during cue presentation. (M) Maximum peaks of reward retrieval onset during CS+ and CS-.

### Stimulation of VGluT3 neurons in the anteromedial BNST has homeostatic consequences

Based on larger BNST VGluT3 neuronal activity during sated sucrose consumption compared with fasted sucrose consumption, we hypothesized that imposing VGluT3 neuronal activation on fasted sucrose consumption would reduce intake. VGluT3-IRES::Cre mice were injected with Cre-dependent GFP or Cre-dependent CoChR-GFP and implanted with stimulating optic fibers (**Figure 6A-B**). Mice were habituated to a sucrose consumption environment before undergoing 4 different stimulation conditions: open-loop, closed-loop, prior, and tonic (GFP=8, 4 female, 4 male, CoChR=7, 4 female, 3 male) (**Figure 6C**). There was a significant effect of stimulation condition on the sucrose ratio, which was calculated as the number of licks when stimulation was on divided by the number of licks when stimulation was off (F(3, 52)=19.6, p<0.00001) (**Figure 6D**). Bonferonni-corrected pairwise comparisons revealed a significant reduction in sucrose ratio during CoChR stimulation, which was not observed in GFP mice, in response to open-loop stimulation (t(7.21)=2.43, p=0.04, CoChR=0.39 ± 0.05, GFP=0.78 ± 0.44), closed-loop stimulation (t(12.9)=2.81, p=0.01, CoChR=0.56 ± 0.15, GFP=0.81 ± 0.18), and tonic stimulation (t(11.7)=3.69, p=0.003, CoChR=0.77 ± 0.18, GFP=1.23 ± 0.29). No difference was found between GFP and CoChR mice in response to prior stimulation, indicating that BNST VGluT3 stimulation reduces sucrose intake only when concurrent with consumption.

**FIGURE 6.**
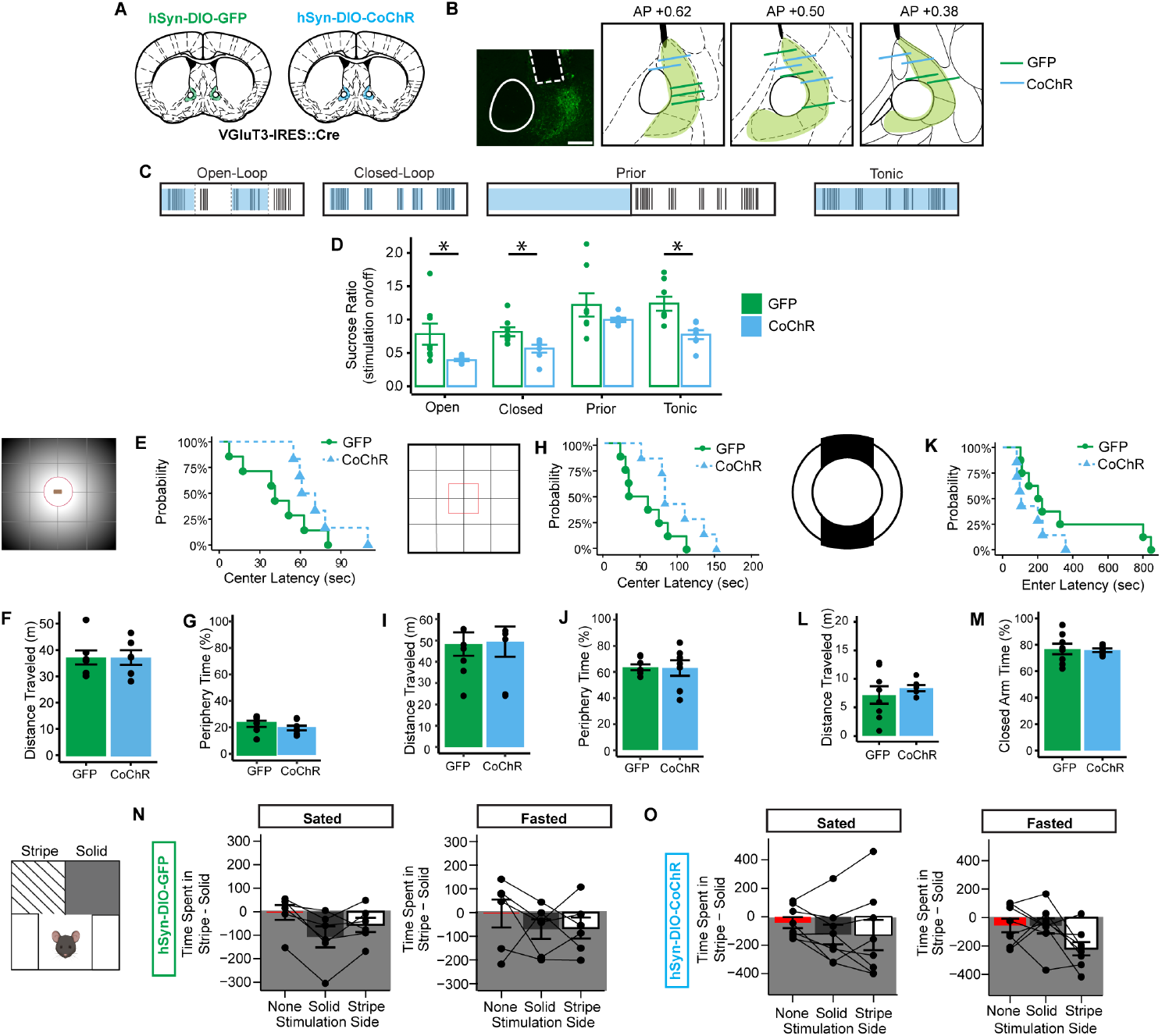
Activation of VGluT3 BNST neurons rapidly reduced sucrose intake in a fasted state but had no effect on anxiety-like behavior nor had intrinsic rewarding or aversive properties. (A) VGluT3-IRES::Cre mice were injected with either Cre-dependent GFP or CoChR. (B) Fiber placement was verified through histology. Scale bar is 200 μm. (C) Illustration of the four stimulation conditions: open-loop, closed-loop, prior, and tonic. (D) The effect of four stimulation conditions on sucrose consumption. GFP=8 (4 female, 4 male), CoChR=7 (4 female, 3 male). Tonic 20Hz optogenetic stimulation during novelty-suppressed feeding on (E) latency to approach the center, (F) distance traveled, and (G) periphery time. Tonic 20Hz optogenetic stimulation during open field when GFP mice (N=8, 4 female, 4 male) and CoChR mice (N=7, 4 female, 3 male) were fasted on (H) latency to approach the center, (I) distance traveled, and (J) periphery time. Tonic 20Hz optogenetic stimulation during elevated zero maze when GFP mice (N=8, 4 female, 4 male) and CoChR mice (N=7, 4 female, 3 male) were fasted on (K) latency to enter the opposite closed arm, (L) distance traveled, and (M) closed arm time. (N) GFP mice (N=6, 3 female, 3 male) in real-time place preference test when sated and fasted. (O) CoChR mice (N=8, 4 female, 4 male) in real-time place preference test when sated and fasted.

Given the BNST is involved in the expression of anxiety-like behavior(24-36), we explored the role of VGluT3 BNST activation in classic anxiety-like behavior paradigms. The novelty-suppressed feeding task evaluates rodent approach-avoidance behavior (64). In this task, 24-hour food deprivation acts as the source of motivation for approach, while the novel environment of an open field and bright light over the food source acts as the source for avoidance. Tonic optogenetic stimulation of GFP mice (N=7, 4 female, 3 male) and CoChR mice (N=6, 3 female, 3 male) during novelty-suppressed feeding had no effect on latency to approach the center (**Figure 6E**), distance traveled (**Figure 6F**), and time spent in the periphery (**Figure 6G**). The novelty-suppressed feeding task necessitates that subjects be fasted, so for appropriate comparison, mice were also fasted when tested in the open field and elevated zero maze, which are also traditional tasks for measuring anxiety-like behavior in rodents. Tonic stimulation for GFP mice (N=8, 4 female, 4 male) and CoChR mice (N=7, 4 female, 3 male) during open field had no group differences in latency to approach the center (**Figure 6H**), distance traveled (**Figure 6I**), and periphery time (**Figure 6J**). There were also no group differences between GFP (N=8, 4 female, 4 male) and CoChR (N=7, 4 female, 3 male) during elevated zero maze in latency to enter the other closed arm (**Figure 6K**), distance traveled (**Figure 6L)**, and closed-arm time (**Figure 6M**). Finally, the valence of VGluT3 BNST activation when sated or fasted was assessed using the real-time place preference task. As expected, GFP mice (N=6, 3 female, 3 male) showed neither preference nor aversion to optogenetic stimulation when sated or fasted. Similarly, CoChR mice (N=8, 4 female, 4 male) showed no preference or aversion to stimulation when sated or fasted. Taken together, VGluT3 BNST neuron activation decreases feeding when fasted and this effect is not driven by aversion, anxiety-like behavior, a or competing reward process.

## DISCUSSION

The BNST is a cellularly diverse brain region linked with anxiety and stress disorders in humans(33, 65-71), as well as feeding(17-23), stress(70-75), and anxiety(24-36) in animal models. Here, we identified a unique BNST cell-type that transcriptionally co-expresses VGluT3 and VGaT. In spite of their transcriptional identity, BNST VGluT3 neurons functionally transmit GABA to the ARC and PVN with rare glutamate co-transmission to the ARC. BNST VGluT3 neurons are preferentially activated by sucrose consumption in the sated state compared with the fasted state and their activation reduces sucrose intake in the fasted state without affecting anxiety-like behavior nor resulting in rewarding or aversive processing.

BNST VGluT3 neurons were located in a distinct location within the anteromedial BNST, part of the extended amygdala. The term “extended amygdala” was derived from observations made through anterograde tract tracing methods that displayed a high degree of contiguity between the central amygdala, medial amygdala, and BNST(1, 76-81). In the same tract tracing studies, the caudomedial end of the nucleus accumbens shell merges with the rostral portion of the BNST(76, 81-83). VGluT3 neurons were observed within the caudomedial shell at low concentration and within the anteromedial BNST VGluT3 neurons were highly concentrated and more lateral compared with the caudomedial shell of the nucleus accumbens(84). In addition, striatal VGluT3 neurons, including those within the posterior nucleus accumbens shell, co-expressed ChAT immunolabeling(44, 56), while VGluT3 BNST neurons did not. Thus, VGluT3 BNST neurons are distinct from striatal VGluT3 neurons.

Our *in situ* hybridization data indicated that VGluT3 BNST neurons largely co-expressed VGluT3 and VGaT mRNA, extending earlier descriptions of VGluT3 mRNA in BNST(38, 85-87). The VGluT3 eurons were a small fraction of the total VGaT-expressing BNST population, indicating they are a unique subset of BNST GABA neurons. As a genetic homologue, VGluT3 shares ∼70% similarity to VGluT1 and VGluT2 and functions as a proton-dependent vesicular glutamate transporter(87-89). Substrate selectivity for glutamate and not other neurotransmitters, including GABA, acetylcholine, and serotonin, was confirmed with its discovery(87-89). VGluT3 is often co-expressed in neurons that co-transmit other neurotransmitters, such as in striatal cholinergic interneurons, raphe serotonin neurons, or retinal glycine neurons(86-91). Thus, genetic evidence suggests VGluT3 BNST neurons could be capable of co-releasing glutamate and GABA but cannot be determined without functional validation through whole-cell electrophysiology.

Before testing whether VGluT3 BNST neurons co-transmit glutamate and GABA, we determined the projection targets of these neurons using Cre-dependent labeling of synaptophysin in VGluT3 BNST processes(57). Consistent with results from PHA-L tracing studies from the anteromedial BNST(92), VGluT3 neurons in the anteromedial BNST projected to the nucleus accumbens, PVN, ARC, and VTA. We also found extensive mRuby expression throughout the BNST, both anterior and posterior, indicating that BNST VGluT3 neurons provide significant input within the BNST. While PHA-L tracing demonstrates that anteromedial BNST neurons also project to the central amygdala, supramammillary and tuberomammillary nuclei, and ventrolateral periaqueductal gray(92), we found no projections to these locations from VGluT3 neurons, indicating a selective projections from this unique cell-type. Presumably other anteromedial BNST cell-types synapse within the central amygdala, mammillary nuclei, and the periaqueductal gray. While synaptophysin labeling was observed in the VTA, similar to previous neuroanatomy studies, the labeling from the anteromedial BNST was minor in comparison to their other targets(93). We immunolabeled for VGluT3 and VGaT protein in the two most prominent targets of VGluT3 BNST neurons: ARC and PVN. In both ARC and PVN, we found that there was approximately 50% co-localization of VGaT out of total VGluT3 protein expression sites.

After identifying the ARC and PVN as primary targets of BNST VGluT3 neurons, we next explored the synaptic functionality of BNST VGluT3 projections by optogenetic terminal stimulation and whole-cell patch-clamp recordings in the ARC and PVN. We found that while VGluT3 BNST neurons have the capability to accumulate glutamate into synaptic vesicles, they functionally transmit GABA, and a small proportion of projections co-transmit glutamate in the ARC. In contrast to the ARC, VGluT3 BNST neurons functionally transmit GABA alone to PVN, as we found no evidence that these neurons transmit glutamate in PVN. These results suggest similar but not identical mechanisms of neurotransmission from VGluT3 BNST neurons to hypothalamic regions. Indeed, neurotransmission from other VGluT3-expressing neurons is not equal across brain regions, with others documenting both GABA and glutamate co-transmission(94, 95) or only glutamate transmission(96, 97). Our findings corroborate previous electron microscopy studies that revealed VGluT3-expressing terminals formed some asymmetrical synapses, but a large majority also formed symmetrical synapses, suggestive of inhibitory neurotransmission(86, 89). Another study also found a large proportion of VGaT protein expression that was absent of VGluT3 protein expression from BNST to VTA(85), suggesting the BNST VGluT3 pathway to VTA also transmits GABA. A growing body of literature suggests that VGluT3 has a synergistic effect, increasing rate of vesicular filling of non-glutamate neurotransmitters, which is driven by the co-expression of VGluT3 and the secondary transporter on the same vesicles, such as acetylcholine(56, 98-100). Whether VGluT3 shows vesicular synergy with VGaT in BNST hypothalamic projections requires further investigation.

Based on the BNST VGluT3 projections to ARC and PVN that participate in homeostatic regulation, we used in vivo fiber photometry to observe calcium-dependent signaling of VGluT3 neurons to rewarding and aversive stimuli during fasted and sated states. BNST VGluT3 neurons were highly activated by aversive footshock stimuli and these signals did not differ between fasted versus sated states. However, VGluT3 neuronal activity was significantly higher in response to sucrose consumption when sated compared to when the mice consumed sucrose when fasted. This signaling pattern is different from the mesolimbic dopamine system where VTA and accumbens neurons show higher activation following reward consumption in the fasted state compared with the sated state(101-104). We also assessed whether BNST VGluT3 neurons were sensitive to Pavlovian predictors of sucrose delivery or non-delivery in the fasted state and found no modulation by the CS+ and CS-. Consistent with free sucrose data in the fasted state, there was also no significant difference in signaling from baseline when the mice entered the magazine to consume the reward during CS+ or CS-. Thus, VGluT3 BNST neurons do not appear to be involved in associative reward learning in the fasted state. Together, VGluT3 neuronal activity is sensitive to internal state in response to a palatable reward but is sensitive to aversive stimuli regardless of internal state.

Based on BNST VGluT3 neuron signaling of state-dependent feeding as well as aversive stimuli, we tested whether optogenetic activation of BNST VGluT3 neurons results in changes in feeding or anxiety-like behavior. We found that optogenetic activation of BNST VGluT3 neurons resulted in a specific behavioral functionality on feeding but not anxiety-like behavior. Activation of VGluT3 BNST neurons decreased feeding in a fasted state but had no effect on classical anxiety-like behavior paradigms: novelty-suppressed feeding, open field, and elevated zero maze. Additionally, stimulation prior to palatable reward access had no effect on consumption, indicating that BNST VGluT3 activation must be concurrent with consumption in the fasted state to reduce feeding. The activation of VGluT3 BNST neurons also had no innate rewarding or aversive valence as assessed in the real-time place conditioning task. Thus, VGluT3 BNST neurons have a specific role in feeding without affecting rewarding, aversive, or anxiety-like behaviors. Given the larger role of BNST in both feeding and anxiety-like behavior, the role of BNST cell-types in feeding and anxiety-like behavior may be better visualized as a Venn diagram with some cell-types distinctly involved in feeding but not anxiety, others in both, and another category that is involved in anxiety but not feeding.

Like the BNST itself, the ARC is diverse in cell-types and not uniform in neurotransmitter release(105-107). The two most well studied cell-types in the ARC are agouti-related peptide (AgRP) and pro-opiomelanocortin (POMC) neurons(108). Classically, the activity of AgRP neurons increases food intake, while activity of POMC neurons decreases food intake(61, 62, 109-111). Activation of POMC neurons also increase heart rate and blood glucose(112, 113). Based on neuroanatomical retrograde tracing, the BNST equally provides input to POMC and AgRP neurons(114). With our combined observations of a reduction in feeding by VGluT3 BNST activation and a largely inhibitory input to the ARC, we hypothesize that VGluT3 BNST neurons preferentially project to AgRP neurons.

The PVN is a critical node in the HPA-axis and plays a central role in stress regulation (63). The PVN is divided into the parvocellular and magnocellular divisions (115, 116), and unbiased transcriptional classification have further revealed that specific cell-types are more populated in discrete subregions of the PVN. For example, Gad2, a genetic marker of GABA synthesis, is comparatively more enriched in the anterior PVN than either the medial or posterior PVN (117). Based on our synaptophysin tracing of BNST VGluT3 efferents, we hypothesize that VGluT3 BNST neurons project to the anterior PVN, as indicated by the oval-shape expression rather than the heart-shape of the posterior PVN. Like the ARC, the phenotypic identity of PVN neurons receiving input from VGluT3 BNST neurons will require further investigation.

In conclusion, we identified a unique cell-type in the anteromedial BNST that transcriptionally co-expresses VGluT3 and VGaT mRNA. VGluT3 BNST neurons transmit GABA in the PVN and ARC with rare glutamate co-transmission in the ARC. These neurons are activated by aversive stimuli and show a state-dependent activation following sucrose consumption in the sated state and not in the fasted state. Activation of VGluT3 BNST neurons reduces sucrose intake in the fasted state and does not affect anxiety-like behavior or innate rewarding or aversive processing. The role of this newly-defined BNST input to the ARC and PVN may prove to be important in understanding the neural pathways involved in energy balance and feeding regulation.

## ACKNOWLEDGEMENTS

This research was supported by the Webb-Waring Biomedical Research Award from the Boettcher Foundation (DHR), Institute for Cannabis Research (DHR),AB Nexus from University of Colorado’s Boulder campus and Anschutz Medical Campus (DHR/CPF), National Institute on Drug Abuse DA047443 (DHR) & DA35821 (CPF), the National Institute on Mental Health MH13222 (AL), and the National Institute of Neurological Disorders and Stroke NS95809 (CPF). AL, DHR, and CPF contributed to experimental design and manuscript writing. AL, DHR, CPF, and RLS contributed to data interpretation. AL, DHR, RK, EDP, HH, and JP contributed to data collection. All authors edited and reviewed the manuscript for publication.

## REFERENCES

1. Alheid GF (2003): Extended Amygdala and Basal Forebrain. Annals of the New York Academy of Sciences. 985:185–205.

2. Bota M, Sporns O, Swanson LW (2012): Neuroinformatics analysis of molecular expression patterns and neuron populations in gray matter regions: The rat BST as a rich exemplar. Brain Research. 1450:174–193.

3. Ju G, Swanson LW, Simerly RB (1989): Studies on the cellular architecture of the bed nuclei of the stria terminalis in the rat: II. chemoarchitecture. Journal of Comparative Neurology. 280:603–621.

4. Ju G, Swanson LW (1989): Studies on the cellular architecture of the bed nuclei of the stria terminalis in the rat: I. cytoarchitecture. Journal of Comparative Neurology. 280:587–602.

5. Dong H-W, Swanson LW (2006): Projections from bed nuclei of the stria terminalis, magnocellular nucleus: Implications for cerebral hemisphere regulation of micturition, defecation, and penile erection. Journal of Comparative Neurology. 494:108–141.

6. Dong H-W, Swanson LW (2006): Projections from bed nuclei of the stria terminalis, anteromedial area: Cerebral hemisphere integration of neuroendocrine, autonomic, and behavioral aspects of energy balance. Journal of Comparative Neurology. 494:142–178.

7. Dong H-W, Swanson LW (2006): Projections from bed nuclei of the stria terminalis, dorsomedial nucleus: Implications for cerebral hemisphere integration of neuroendocrine, autonomic, and drinking responses. Journal of Comparative Neurology. 494:75–107.

8. Dong H-W, Swanson LW (2004): Projections from bed nuclei of the stria terminalis, posterior division: Implications for cerebral hemisphere regulation of defensive and reproductive behaviors. Journal of Comparative Neurology. 471:396–433.

9. Dong H-W, Swanson LW (2004): Organization of axonal projections from the anterolateral area of the bed nuclei of the stria terminalis. Journal of Comparative Neurology. 468:277–298.

10. Dong H-W, Swanson LW (2003): Projections from the rhomboid nucleus of the bed nuclei of the stria terminalis: Implications for cerebral hemisphere regulation of ingestive behaviors. Journal of Comparative Neurology. 463:434–472.

11. Dong HW, Petrovich GD, Swanson LW (2000): Organization of projections from the juxtacapsular nucleus of the BST: a PHAL study in the rat. Brain Research. 859:1–14.

12. Dong H-W, Petrovich GD, Watts AG, Swanson LW (2001): Basic organization of projections from the oval and fusiform nuclei of the bed nuclei of the stria terminalis in adult rat brain. Journal of Comparative Neurology. 436:430–455.

13. Krettek JE, Price JL (1978): Amygdaloid projections to subcortical structures within the basal forebrain and brainstem in the rat and cat. J Comp Neurol. 178:225–254.

14. Gungor NZ, Pare D (2016): Functional Heterogeneity in the Bed Nucleus of the Stria Terminalis. J Neurosci. 36:8038–8049.

15. Moga MM, Saper CB (1994): Neuropeptide-immunoreactive neurons projecting to the paraventricular hypothalamic nucleus in the rat. J Comp Neurol. 346:137–150.

16. Barbier M, González JA, Houdayer C, Burdakov D, Risold PY, Croizier S (2021): Projections from the dorsomedial division of the bed nucleus of the stria terminalis to hypothalamic nuclei in the mouse. J Comp Neurol. 529:929–956.

17. Zhang J, Wang L, Yang Y, Wang S, Huang C, Yang L, et al. (2023): Dissection of the bed nucleus of the stria terminalis neuronal subtypes in feeding regulation. Physiol Behav. 271:114333.

18. de Araujo Salgado I, Li C, Burnett CJ, Rodriguez Gonzalez S, Becker JJ, Horvath A, et al. (2023): Toggling between food-seeking and self-preservation behaviors via hypothalamic response networks. Neuron.

19. Luskin AT, Bhatti DL, Mulvey B, Pedersen CE, Girven KS, Oden-Brunson H, et al. (2021): Extended amygdala-parabrachial circuits alter threat assessment and regulate feeding. Science Advances. 7:eabd3666.

20. Jennings JH, Rizzi G, Stamatakis AM, Ung RL, Stuber GD (2013): The Inhibitory Circuit Architecture of the Lateral Hypothalamus Orchestrates Feeding. Science. 341:1517–1521.

21. Jaramillo AA, Williford KM, Marshall C, Winder DG, Centanni SW (2020): BNST transient activity associates with approach behavior in a stressful environment and is modulated by the parabrachial nucleus. Neurobiology of Stress. 13:100247.

22. Jia X, Chen S, Li X, Tao S, Lai J, Liu H, et al. (2022): Divergent neurocircuitry dissociates two components of the stress response: glucose mobilization and anxiety-like behavior. Cell Reports. 41:111586.

23. Douglass AM, Resch JM, Madara JC, Kucukdereli H, Yizhar O, Grama A, et al. (2023): Neural basis for fasting activation of the hypothalamic–pituitary–adrenal axis. Nature. 620:154–162.

24. Fendt M, Endres T, Apfelbach R (2003): Temporary Inactivation of the Bed Nucleus of the Stria Terminalis But Not of the Amygdala Blocks Freezing Induced by Trimethylthiazoline, a Component of Fox Feces. The Journal of Neuroscience. 23:23–28.

25. Sullivan GM, Apergis J, Bush DE, Johnson LR, Hou M, Ledoux JE (2004): Lesions in the bed nucleus of the stria terminalis disrupt corticosterone and freezing responses elicited by a contextual but not by a specific cue-conditioned fear stimulus. Neuroscience. 128:7–14.

26. Waddell J, Morris RW, Bouton ME (2006): Effects of bed nucleus of the stria terminalis lesions on conditioned anxiety: Aversive conditioning with long-duration conditional stimuli and reinstatement of extinguished fear. Behavioral Neuroscience. 120:324–336.

27. Sajdyk T, Johnson P, Fitz S, Shekhar A (2008): Chronic inhibition of GABA synthesis in the bed nucleus of the stria terminalis elicits anxiety-like behavior. Journal of Psychopharmacology. 22:633–641.

28. Duvarci S, Bauer EP, Paré D (2009): The Bed Nucleus of the Stria Terminalis Mediates Inter-individual Variations in Anxiety and Fear. The Journal of Neuroscience. 29:10357–10361.

29. Hammack SE, Cheung J, Rhodes KM, Schutz KC, Falls WA, Braas KM, et al. (2009): Chronic stress increases pituitary adenylate cyclase-activating peptide (PACAP) and brain-derived neurotrophic factor (BDNF) mRNA expression in the bed nucleus of the stria terminalis (BNST): roles for PACAP in anxiety-like behavior. Psychoneuroendocrinology. 34:833–843.

30. Davis M, Walker DL, Miles L, Grillon C (2010): Phasic vs Sustained Fear in Rats and Humans: Role of the Extended Amygdala in Fear vs Anxiety. Neuropsychopharmacology. 35:105–135.

31. Ventura-Silva AP, Pêgo JM, Sousa JC, Marques AR, Rodrigues AJ, Marques F, et al. (2012): Stress shifts the response of the bed nucleus of the stria terminalis to an anxiogenic mode. Eur J Neurosci. 36:3396–3406.

32. Breitfeld T, Bruning J, Inagaki H, Takeuchi Y, Kiyokawa Y, Fendt M (2015): Temporary inactivation of the anterior part of the bed nucleus of the stria terminalis blocks alarm pheromone-induced defensive behavior in rats. Frontiers in Neuroscience. 9.

33. Buff C, Brinkmann L, Bruchmann M, Becker MPI, Tupak S, Herrmann MJ, et al. (2017): Activity alterations in the bed nucleus of the stria terminalis and amygdala during threat anticipation in generalized anxiety disorder. Social Cognitive and Affective Neuroscience. 12:1766–1774.

34. Mazzone CM, Pati D, Michaelides M, DiBerto J, Fox JH, Tipton G, et al. (2018): Acute engagement of Gq-mediated signaling in the bed nucleus of the stria terminalis induces anxiety-like behavior. Molecular Psychiatry. 23:143–153.

35. Yamauchi N, Takahashi D, Sugimura YK, Kato F, Amano T, Minami M (2018): Activation of the neural pathway from the dorsolateral bed nucleus of the stria terminalis to the central amygdala induces anxiety-like behaviors. European Journal of Neuroscience. 48:3052–3061.

36. Williford KM, Taylor A, Melchior JR, Yoon HJ, Sale E, Negasi MD, et al. (2023): BNST PKCδ neurons are activated by specific aversive conditions to promote anxiety-like behavior. Neuropsychopharmacology. 48:1031–1041.

37. Siletti K, Hodge R, Mossi Albiach A, Hu L, Lee KW, Lönnerberg P, et al. (2022): Transcriptomic diversity of cell types across the adult human brain. bioRxiv.2022.2010.2012.511898.

38. Welch JD, Kozareva V, Ferreira A, Vanderburg C, Martin C, Macosko EZ (2019): Single-Cell Multi-omic Integration Compares and Contrasts Features of Brain Cell Identity. Cell. 177:1873–1887 e1817.

39. Giardino WJ, Eban-Rothschild A, Christoffel DJ, Li S-B, Malenka RC, de Lecea L (2018): Parallel circuits from the bed nuclei of stria terminalis to the lateral hypothalamus drive opposing emotional states. Nature Neuroscience. 21:1084–1095.

40. Kim S-Y, Adhikari A, Lee SY, Marshel JH, Kim CK, Mallory CS, et al. (2013): Diverging neural pathways assemble a behavioural state from separable features in anxiety. Nature. 496:219–223.

41. Jennings JH, Sparta DR, Stamatakis AM, Ung RL, Pleil KE, Kash TL, et al. (2013): Distinct extended amygdala circuits for divergent motivational states. Nature. 496:224–228.

42. Han RW, Zhang ZY, Jiao C, Hu ZY, Pan BX (2024): Synergism between two BLA-to-BNST pathways for appropriate expression of anxiety-like behaviors in male mice. Nat Commun. 15:3455.

43. Fremeau RT, Jr., Burman J, Qureshi T, Tran CH, Proctor J, Johnson J, et al. (2002): The identification of vesicular glutamate transporter 3 suggests novel modes of signaling by glutamate. Proc Natl Acad Sci U S 99:14488–14493.

44. Gras C, Herzog E, Bellenchi GC, Bernard V, Ravassard P, Pohl M, et al. (2002): A third vesicular glutamate transporter expressed by cholinergic and serotoninergic neurons. J Neurosci. 22:5442–5451.

45. Schafer MK, Varoqui H, Defamie N, Weihe E, Erickson JD (2002): Molecular cloning and functional identification of mouse vesicular glutamate transporter 3 and its expression in subsets of novel excitatory neurons. J Biol Chem. 277:50734–50748.

46. Franklin KBJ, Paxinos G (2013): Paxinos and Franklin’s The mouse brain in stereotaxic coordinates. Fourth edition. ed. Amsterdam: Academic Press, an imprint of Elsevier.

47. Yang Y, Atasoy D, Su HH, Sternson SM (2011): Hunger states switch a flip-flop memory circuit via a synaptic AMPK-dependent positive feedback loop. Cell. 146:992–1003.

48. Hentges ST, Nishiyama M, Overstreet LS, Stenzel-Poore M, Williams JT, Low MJ (2004): GABA Release from Proopiomelanocortin Neurons. The Journal of Neuroscience. 24:1578–1583.

49. Luther JA, Tasker JG (2000): Voltage-gated currents distinguish parvocellular from magnocellular neurones in the rat hypothalamic paraventricular nucleus. The Journal of Physiology. 523:193–209.

50. Barker DJ, Miranda-Barrientos J, Zhang S, Root DH, Wang HL, Liu B, et al. (2017): Lateral Preoptic Control of the Lateral Habenula through Convergent Glutamate and GABA Transmission. Cell Rep. 21:1757–1769.

51. McGovern DJ, Ly A, Ecton KL, Huynh DT, Prévost ED, Gonzalez SC, et al. (2024): Ventral tegmental area glutamate neurons mediate nonassociative consequences of stress. Mol Psychiatry. 29:1671–1682.

52. McGovern DJ, Polter AM, Root DH (2021): Neurochemical Signaling of Reward and Aversion to Ventral Tegmental Area Glutamate Neurons. The Journal of Neuroscience. 41:5471–5486.

53. Aitken TJ, Ly T, Shehata S, Sivakumar N, Medina NS, Gray LA, et al. (2023): Negative feedback control of hunger circuits by the taste of food. bioRxiv.

54. Chen Y, Lin Y-C, Zimmerman CA, Essner RA, Knight ZA (2016): Hunger neurons drive feeding through a sustained, positive reinforcement signal. eLife. 5:e18640.

55. O’Connor Eoin C, Kremer Y, Lefort S, Harada M, Pascoli V, Rohner C, et al. (2015): Accumbal D1R Neurons Projecting to Lateral Hypothalamus Authorize Feeding. Neuron. 88:553–564.

56. Gras C, Amilhon B, Lepicard ÈM, Poirel O, Vinatier J, Herbin M, et al. (2008): The vesicular glutamate transporter VGLUT3 synergizes striatal acetylcholine tone. Nature Neuroscience. 11:292–300.

57. Beier KT, Steinberg EE, DeLoach KE, Xie S, Miyamichi K, Schwarz L, et al. (2015): Circuit Architecture of VTA Dopamine Neurons Revealed by Systematic Input-Output Mapping. Cell. 162:622–634.

58. Root DH, Barker DJ, Estrin DJ, Miranda-Barrientos JA, Liu B, Zhang S, et al. (2020): Distinct Signaling by Ventral Tegmental Area Glutamate, GABA, and Combinatorial Glutamate-GABA Neurons in Motivated Behavior. Cell Rep. 32:108094.

59. Klapoetke NC, Murata Y, Kim SS, Pulver SR, Birdsey-Benson A, Cho YK, et al. (2014): Independent optical excitation of distinct neural populations. Nature Methods. 11:338–346.

60. Shemesh OA, Tanese D, Zampini V, Linghu C, Piatkevich K, Ronzitti E, et al. (2017): Temporally precise single-cell-resolution optogenetics. Nat Neurosci. 20:1796–1806.

61. Krashes MJ, Koda S, Ye C, Rogan SC, Adams AC, Cusher DS, et al. (2011): Rapid, reversible activation of AgRP neurons drives feeding behavior in mice. J Clin Invest. 121:1424–1428.

62. Aponte Y, Atasoy D, Sternson SM (2011): AGRP neurons are sufficient to orchestrate feeding behavior rapidly and without training. Nature Neuroscience. 14:351–355.

63. Spencer RL, Deak T (2017): A users guide to HPA axis research. Physiol Behav. 178:43–65.

64. Samuels BA, Hen R (2011): Novelty-Suppressed Feeding in the Mouse. In: Gould TD, editor. Mood and Anxiety Related Phenotypes in Mice: Characterization Using Behavioral Tests, Volume II. Totowa, NJ: Humana Press, pp 107–121.

65. Petranu K, Webb EK, Tomas CW, Harb F, Torres L, deRoon-Cassini TA, et al. (2024): Investigating the bed nucleus of the stria terminalis as a predictor of posttraumatic stress disorder in Black Americans and the moderating effects of racial discrimination. Transl Psychiatry. 14:337.

66. Feola B, Flook EA, Gardner H, Phan KL, Gwirtsman H, Olatunji B, et al. (2023): Altered bed nucleus of the stria terminalis and amygdala responses to threat in combat veterans with posttraumatic stress disorder. J Trauma Stress. 36:359–372.

67. Mobbs D, Yu R, Rowe JB, Eich H, FeldmanHall O, Dalgleish T (2010): Neural activity associated with monitoring the oscillating threat value of a tarantula. Proc Natl Acad Sci U S A. 107:20582–20586.

68. Brinkmann L, Buff C, Neumeister P, Tupak SV, Becker MP, Herrmann MJ, et al. (2017): Dissociation between amygdala and bed nucleus of the stria terminalis during threat anticipation in female post-traumatic stress disorder patients. Hum Brain Mapp. 38:2190–2205.

69. Awasthi S, Pan H, LeDoux JE, Cloitre M, Altemus M, McEwen B, et al. (2020): The bed nucleus of the stria terminalis and functionally linked neurocircuitry modulate emotion processing and HPA axis dysfunction in posttraumatic stress disorder. NeuroImage: Clinical. 28:102442.

70. Hu P, Liu J, Maita I, Kwok C, Gu E, Gergues MM, et al. (2020): Chronic Stress Induces Maladaptive Behaviors by Activating Corticotropin-Releasing Hormone Signaling in the Mouse Oval Bed Nucleus of the Stria Terminalis. The Journal of Neuroscience. 40:2519–2537.

71. Maita I, Roepke TA, Samuels BA (2022): Chronic stress-induced synaptic changes to corticotropin-releasing factor-signaling in the bed nucleus of the stria terminalis. Front Behav Neurosci. 16:903782.

72. Hammack SE, Richey KJ, Watkins LR, Maier SF (2004): Chemical lesion of the bed nucleus of the stria terminalis blocks the behavioral consequences of uncontrollable stress. Behav Neurosci. 118:443–448.

73. Roman CW, Lezak KR, Kocho-Schellenberg M, Garret MA, Braas K, May V, et al. (2012): Excitotoxic lesions of the bed nucleus of the stria terminalis (BNST) attenuate the effects of repeated stress on weight gain: Evidence for the recruitment of BNST activity by repeated, but not acute, stress. Behavioural Brain Research. 227:300–304.

74. Zelikowsky M, Hui M, Karigo T, Choe A, Yang B, Blanco MR, et al. (2018): The Neuropeptide Tac2 Controls a Distributed Brain State Induced by Chronic Social Isolation Stress. Cell. 173:1265-1279.e1219.

75. Radley JJ, Sawchenko PE (2015): Evidence for involvement of a limbic paraventricular hypothalamic inhibitory network in hypothalamic-pituitary-adrenal axis adaptations to repeated stress. Journal of Comparative Neurology. 523:2769–2787.

76. Alheid GF, Heimer L (1988): New perspectives in basal forebrain organization of special relevance for neuropsychiatric disorders: The striatopallidal, amygdaloid, and corticopetal components of substantia innominata. Neuroscience. 27:1–39.

77. de Olmos JS, Heimer L (1999): The concepts of the ventral striatopallidal system and extended amygdala. Ann N Y Acad Sci. 877:1–32.

78. De Olmos JS, Ingram WR (1972): The projection field of the stria terminalis in the rat brain. An experimental study. J Comp Neurol. 146:303–334.

79. Dong HW, Petrovich GD, Swanson LW (2001): Topography of projections from amygdala to bed nuclei of the stria terminalis. Brain Res Brain Res Rev. 38:192–246.

80. Weller KL, Smith DA (1982): Afferent connections to the bed nucleus of the stria terminalis. Brain Res. 232:255–270.

81. de Olmos S, Lorenzo A (2023): Developing the theory of the extended amygdala with the use of the cupric-silver technique. Journal of the History of the Neurosciences. 32:19–38.

82. Woodhams PL, Roberts GW, Polak JM, Crow TJ (1983): Distribution of neuropeptides in the limbic system of the rat: the bed nucleus of the stria terminalis, septum and preoptic area. Neuroscience. 8:677–703.

83. Heimer L, Harlan RE, Alheid GF, Garcia MM, de Olmos J (1997): Substantia innominata: a notion which impedes clinical-anatomical correlations in neuropsychiatric disorders. Neuroscience. 76:957–1006.

84. Zahm DS (1998): Is the caudomedial shell of the nucleus accumbens part of the extended amygdala? A consideration of connections. Crit Rev Neurobiol. 12:245–265.

85. Kudo T, Uchigashima M, Miyazaki T, Konno K, Yamasaki M, Yanagawa Y, et al. (2012): Three Types of Neurochemical Projection from the Bed Nucleus of the Stria Terminalis to the Ventral Tegmental Area in Adult Mice. The Journal of Neuroscience. 32:18035–18046.

86. Fremeau RT, Burman J, Qureshi T, Tran CH, Proctor J, Johnson J, et al. (2002): The identification of vesicular glutamate transporter 3 suggests novel modes of signaling by glutamate. Proceedings of the National Academy of Sciences. 99:14488–14493.

87. Schäfer MKH, Varoqui H, Defamie N, Weihe E, Erickson JD (2002): Molecular Cloning and Functional Identification of Mouse Vesicular Glutamate Transporter 3 and Its Expression in Subsets of Novel Excitatory Neurons. Journal of Biological Chemistry. 277:50734–50748.

88. Takamori S, Malherbe P, Broger C, Jahn R (2002): Molecular cloning and functional characterization of human vesicular glutamate transporter 3. EMBO reports. 3:798-803-803.

89. Gras C, Herzog E, Bellenchi GC, Bernard V, Ravassard P, Pohl M, et al. (2002): A Third Vesicular Glutamate Transporter Expressed by Cholinergic and Serotoninergic Neurons. The Journal of Neuroscience. 22:5442.

90. Haverkamp S, Wässle H (2004): Characterization of an amacrine cell type of the mammalian retina immunoreactive for vesicular glutamate transporter 3. Journal of Comparative Neurology. 468:251–263.

91. Johnson J, Sherry DM, Liu X, Fremeau Jr. RT, Seal RP, Edwards RH, et al. (2004): Vesicular glutamate transporter 3 expression identifies glutamatergic amacrine cells in the rodent retina. Journal of Comparative Neurology. 477:386–398.

92. Dong H-W, Swanson LW (2006): Projections from bed nuclei of the stria terminalis, anteromedial area: cerebral hemisphere integration of neuroendocrine, autonomic, and behavioral aspects of energy balance. The Journal of comparative neurology. 494:142–178.

93. Zahm DS, Williams EA, Latimer MP, Winn P (2001): Ventral mesopontine projections of the caudomedial shell of the nucleus accumbens and extended amygdala in the rat: Double dissociation by organization and development. Journal of Comparative Neurology. 436:111–125.

94. Pelkey KA, Calvigioni D, Fang C, Vargish G, Ekins T, Auville K, et al. (2020): Paradoxical network excitation by glutamate release from VGluT3+ GABAergic interneurons. eLife. 9:e51996.

95. Stinson HE, Ninan I (2024): Median raphe glutamatergic neuron-mediated enhancement of GABAergic transmission and suppression of long-term potentiation in the hippocampus. Heliyon. 10:e38192.

96. Wang HL, Zhang S, Qi J, Wang H, Cachope R, Mejias-Aponte CA, et al. (2019): Dorsal Raphe Dual Serotonin-Glutamate Neurons Drive Reward by Establishing Excitatory Synapses on VTA Mesoaccumbens Dopamine Neurons. Cell Rep. 26:1128–1142 e1127.

97. Qi J, Zhang S, Wang HL, Wang H, de Jesus Aceves Buendia J, Hoffman AF, et al. (2014): A glutamatergic reward input from the dorsal raphe to ventral tegmental area dopamine neurons. Nat Commun. 5:5390.

98. Favier M, Martin Garcia E, Icick R, de Almeida C, Jehl J, Desplanque M, et al. (2024): The human VGLUT3-pT8I mutation elicits uneven striatal DA signaling, food or drug maladaptive consumption in male mice. Nat Commun. 15:5691.

99. Amilhon B, Lepicard E, Renoir T, Mongeau R, Popa D, Poirel O, et al. (2010): VGLUT3 (vesicular glutamate transporter type 3) contribution to the regulation of serotonergic transmission and anxiety. J Neurosci. 30:2198–2210.

100. Nelson AB, Bussert TG, Kreitzer AC, Seal RP (2014): Striatal cholinergic neurotransmission requires VGLUT3. J Neurosci. 34:8772–8777.

101. McGovern DJ, Phillips A, Ly A, Prévost ED, Ward L, Siletti K, et al. (2024): Salience signaling and stimulus scaling of ventral tegmental area glutamate neuron subtypes. bioRxiv.2024.2006. 2012.598688.

102. Liu S, Borgland SL (2015): Regulation of the mesolimbic dopamine circuit by feeding peptides. Neuroscience. 289:19–42.

103. Wallace CW, Loudermilt MC, Fordahl SC (2022): Effect of fasting on dopamine neurotransmission in subregions of the nucleus accumbens in male and female mice. Nutr Neurosci. 25:1338–1349.

104. Hsu TM, McCutcheon JE, Roitman MF (2018): Parallels and Overlap: The Integration of Homeostatic Signals by Mesolimbic Dopamine Neurons. Front Psychiatry. 9:410.

105. Campbell JN, Macosko EZ, Fenselau H, Pers TH, Lyubetskaya A, Tenen D, et al. (2017): A molecular census of arcuate hypothalamus and median eminence cell types. Nature Neuroscience. 20:484–496.

106. Hentges ST, Otero-Corchon V, Pennock RL, King CM, Low MJ (2009): Proopiomelanocortin expression in both GABA and glutamate neurons. J Neurosci. 29:13684–13690.

107. Graebner AK, Iyer M, Carter ME (2015): Understanding how discrete populations of hypothalamic neurons orchestrate complicated behavioral states. Frontiers in systems neuroscience. 9:111.

108. Chen Y, Knight ZA (2016): Making sense of the sensory regulation of hunger neurons. Bioessays. 38:316–324.

109. Luquet S, Perez FA, Hnasko TS, Palmiter RD (2005): NPY/AgRP Neurons Are Essential for Feeding in Adult Mice but Can Be Ablated in Neonates. Science. 310:683–685.

110. Gropp E, Shanabrough M, Borok E, Xu AW, Janoschek R, Buch T, et al. (2005): Agouti-related peptide– expressing neurons are mandatory for feeding. Nature Neuroscience. 8:1289–1291.

111. Qu N, He Y, Wang C, Xu P, Yang Y, Cai X, et al. (2020): A POMC-originated circuit regulates stress-induced hypophagia, depression, and anhedonia. Mol Psychiatry. 25:1006–1021.

112. Üner AG, Keçik O, Quaresma PGF, De Araujo TM, Lee H, Li W, et al. (2019): Role of POMC and AgRP neuronal activities on glycaemia in mice. Sci Rep. 9:13068.

113. Jia X, Chen S, Li X, Tao S, Lai J, Liu H, et al. (2022): Divergent neurocircuitry dissociates two components of the stress response: glucose mobilization and anxiety-like behavior. Cell Rep. 41:111586.

114. Wang D, He X, Zhao Z, Feng Q, Lin R, Sun Y, et al. (2015): Whole-brain mapping of the direct inputs and axonal projections of POMC and AgRP neurons. Front Neuroanat. 9:40.

115. Swanson LW, Sawchenko PE (1983): Hypothalamic Integration: Organization of the Paraventricular and Supraoptic Nuclei. Annual Review of Neuroscience. 6:269–324.

116. Liposits Z (1993): Ultrastructure of hypothalamic paraventricular neurons. Critical reviews in neurobiology. 7:89–162.

117. Xu S, Yang H, Menon V, Lemire AL, Wang L, Henry FE, et al. (2020): Behavioral state coding by molecularly defined paraventricular hypothalamic cell type ensembles. Science. 370.

